# Linking sugar sensing to immunity in plants through O-glycosylation of immune-signaling kinases

**DOI:** 10.64898/2026.05.04.722566

**Authors:** Yalikunjiang Aizezi, Hongliang Zhang, Yang Bi, Jung-Gun Kim, Ajeet Chaudhary, Cao Son Trinh, Qingxi J. Shen, Shou-Ling Xu, Mary Beth Mudgett, Zhi-Yong Wang

## Abstract

Interaction with microbes can reprogram metabolism and alter nutrient availability in plant cells. How metabolic changes modulate immune responses remains unknown. Here, we show that sugar-sensing *O*-glycosylation of immune-signaling kinases mediates metabolic regulation of immunity. Under sugar-replete conditions, the MAP kinase kinases (MKK4 and MKK5), key components of pattern-triggered immunity (PTI), are glycosylated with *O*-GlcNAc and *O*-fucose on their activation loops, blocking their phosphorylation by upstream kinases and thereby restricting PTI. Pathogen infection or sugar starvation reduces *O*-glycosylation of MKK4/5 and enhances immune signaling; these effects are reversed by GDP-fucose treatment, demonstrating that reduced sugar availability decreases *O*-fucosylation and enhances immune signaling in infected cells. Chemical inhibition of *O*-fucosylation enhances immunity and pathogen resistance in both Arabidopsis and tomato. Our findings establish *O*-glycosylation of MKKs as a metabolic rheostat that fine-tunes immune responses according to sugar availability during plant-microbe interactions, providing a new strategy for improving crop health.

## Introduction

Plants interact with diverse microbes that have distinct effects on host metabolism and nutrient status, ranging from beneficial microbes that promote nutrient acquisition to pathogens that cause nutrient depletion. Therefore, controlling immune responses according to metabolic and nutritional changes should be crucial for managing interactions with different microbes.^1^ Furthermore, metabolic regulation of immune intensity is key for effective defense against pathogens while balancing the growth-defense tradeoff and minimizing fitness costs.^1,2^ However, the molecular link between nutrient sensing and immune signaling remains unclear.^1^

Decades of research have established that plants, like other eukaryotes, use pattern-recognition receptors (PRRs) to detect conserved microbial features known as microbe-associated molecular patterns (MAMPs). Recognition of a MAMP initiates immune signaling (known as PTI, or pattern-triggered immunity) and ultimately leads to the production of an array of antimicrobial responses.^3,4^ For example, in *Arabidopsis thaliana*, peptides derived from the bacterial flagella (named flagellin) are perceived by the plasma membrane-associated leucine-rich repeat receptor kinase FLS2 and its co-receptor kinase BAK1.^5^ Activated FLS2/BAK1 receptor complexes initiate a phosphorylation cascade that sequentially activates receptor-like cytoplasmic kinases,^6^ BSU1-family phosphatases,^7^ and MITOGEN-ACTIVATED PROTEIN KINASE (MAPK) modules.^3,8,9^ The MAPK modules include multiple MAP kinase kinase kinases (MAPKKKs/MEKKs), which activate two cascades of MAP kinase kinases (MKKs) and MAP kinases (MPKs): MKK4/MKK5–MPK3/MPK6 and MKK1/MKK2–MPK4.^8–10^ The MPKs phosphorylate various substrate proteins, including transcription factors, channel proteins, and metabolic enzymes, thereby activating defense-related gene expression, antibacterial metabolite biosynthesis, and stomatal immunity.^8,11,12^

However, MAMPs are broadly conserved among microbes, and their recognition by PRRs is insufficient for plants to distinguish pathogens from non-pathogenic microbes and to regulate immune responses appropriately.^1^ A key distinction between pathogenic and non-pathogenic microbes is that pathogens often exploit host nutrients to support their own growth, causing disease.^13^ For example, some bacterial pathogens deploy effector proteins to activate host sugar transporters, such as SWEETs, causing sugar export into the apoplast.^14–16^ Some pathogen effectors alter host metabolism to channel nutrients to the pathogens^17^. Recent studies have shown that fungal pathogens redirect amino acid transporters to sites of infection, thereby extending the concept of transporter-mediated susceptibility.^18^ Consistent with the concept of pathogen exploitation of host nutrients, spatiometabolic analyses have shown reduced levels of sugars and nutrients in infected areas of leaf tissues.^19^ In contrast, beneficial microbes often improve host plants’ nutrient acquisition and suppress their immunity.^1,2^ How the hosts’ immune systems respond to such microbe-induced changes in metabolism and nutrient status is unknown.

*O*-glycosylation of serine and threonine residues on nucleocytoplasmic proteins is an essential sugar-sensing mechanism.^20,21^ In mammals, dynamic addition and removal of *O*-linked *β*−N-acetylglucosamine (*O*-GlcNAc) modifications are catalyzed by a single pair of enzymes, *O*-GlcNAc transferase (OGT) and *O*-GlcNAcase (OGA).^20^ OGT activity depends on the level of the donor substrate UDP-GlcNAc,^22^ which is synthesized through the convergence of primary metabolic pathways and thus reflects cellular nutrient status.^23^ As an essential nutrient-sensing mechanism, *O*-GlcNAc is implicated in numerous human diseases, including neurodegeneration, cancer, diabetes, and immunological disorders.^24,25^ In plants, two OGT homologs, SPINDLY (SPY) and SECRET AGENT (SEC), function as *O*-fucose transferase and *O*-GlcNAc transferase using GDP-fucose and UDP-GlcNAc as donor substrates, respectively.^26,27^ Genetic studies indicate that SPY and SEC are essential for viability and play redundant, antagonistic, or independent roles in various biological processes, including hormone signaling, fruit ripening, circadian rhythm, and stress adaptation.^28,29^ Notably, SPY is essential for seedling growth response to exogenous sugar, supporting its role in sugar sensing.^30^ Proteomic studies identified hundreds of *O*-GlcNAcylated and *O*-fucosylated proteins, predominantly nuclear and cytosolic, with broad signaling and regulatory functions.^30–33^ The proteins modified by *O-*GlcNAc and *O*-fucose include several PTI components.^30^ However, whether *O*-glycosylation mediate metabolic regulation of immune signaling in plant-microbe interactions remains unknown.

Here, we report that sugar status gates immune activation in Arabidopsis through crosstalk between protein *O*-glycosylation and phosphorylation. By combining mass spectrometry, biochemical, and genetic analyses, we show that MAP kinase kinases (MKK4 and MKK5) are *O*-glycosylated with either GlcNAc or fucose at key residues in their activation loops under sugar-replete conditions. These sugar modifications block their phosphorylation by upstream MEKK, thus restricting the magnitude of PTI signaling. By contrast, infection with bacterial pathogen *Pseudomonas syringae* reduces MKK *O*-fucosylation, thereby allowing MKK phosphorylation and enhancing immune signaling. Similarly, chemical inhibition of SPY-mediated *O*-fucosylation enhances MAMP-induced MKK5 phosphorylation and boosts pathogen resistance in Arabidopsis and tomato. Together, our findings establish sugar-dependent *O*-glycosylation of MKKs as a molecular link that mediates metabolic regulation of innate immune signaling in plants.

## Results

### Loss of protein *O*-glycosylation enhances MAMP-triggered immunity

To test whether the glycosylation enzymes SPY and SEC regulate the function of proteins involved in PTI signaling, we first examined how *spy* and *sec* mutant seedlings respond to MAMP elicitation. The flagellin-derived peptide flg22 not only activates PTI but also inhibits seedling root growth. Both *spy-4* and *spy-23* mutants showed hypersensitivity to flg22-induced root growth inhibition, while *sec-2* and *sec-5* showed a comparatively milder hypersensitivity (Figure 1A, B; Figure S1A-B). These mutant phenotypes were restored by complementation – the *spy-23* allele was complemented by a transgene expressing SPY-GFP and the *sec-5* allele was complemented by a *SEC-TurboID* transgene (Figure S1A-B). Consistent with the severity of root growth inhibition, flg22 treatment induced a stronger ROS burst in *spy-4* mutant leaves compared to wild-type and *sec-2* leaves (Figure 1C).

**Figure 1.**
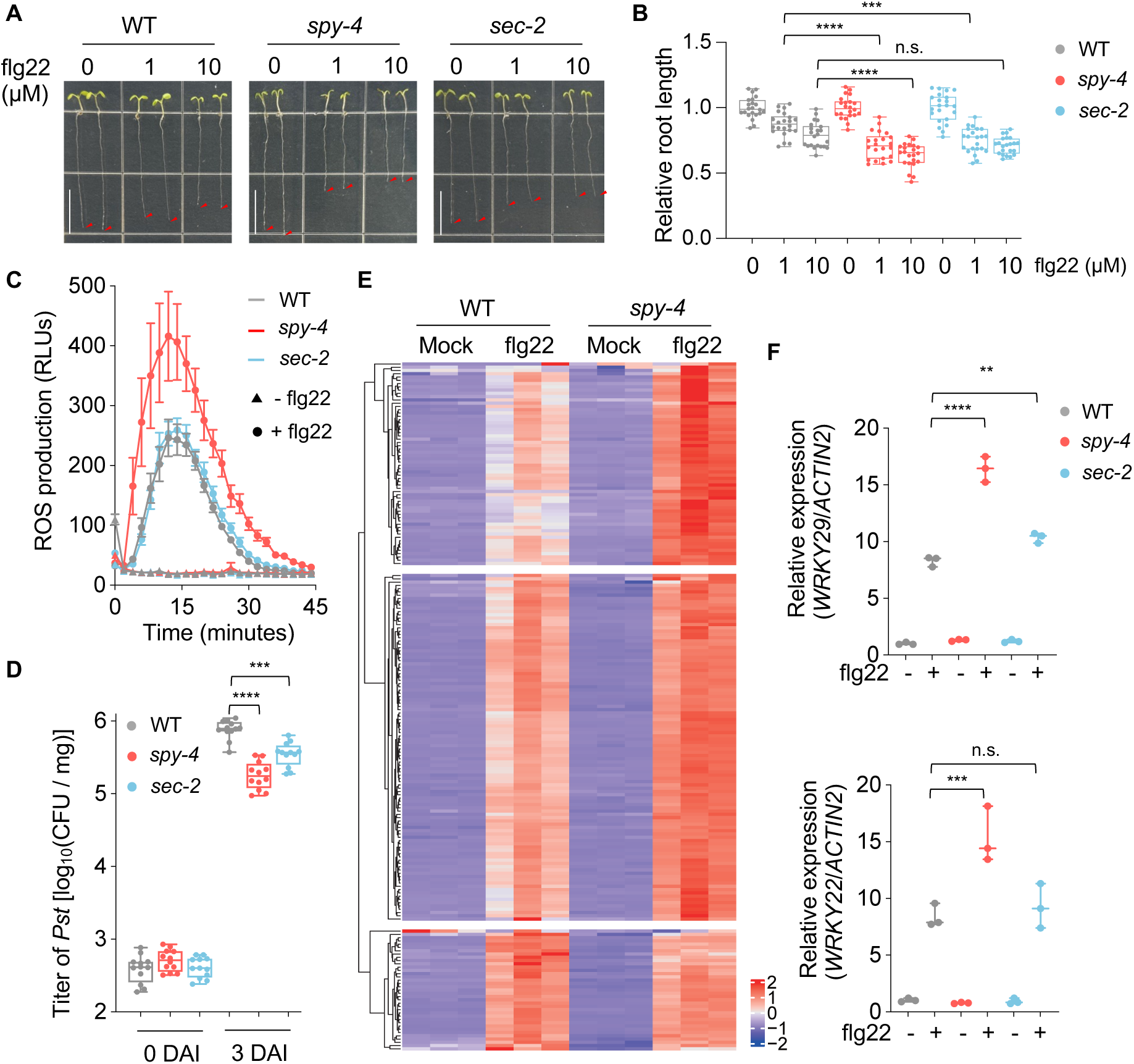
Flg22 and pathogen-induced responses are enhanced in *spy* and *sec* mutants. (A) Primary root length phenotype of 7-day-old wild type (WT), *spy-4*, and *sec-2* mutants grown on ½ MS medium supplemented with or without flg22 at indicated concentrations. Bar = 1 cm. (B) Quantification of seedling root length (n =22 biological replicates) shown in (A). Primary root length was normalized to the mock group. (C) Real-time reading of flg22-induced reactive oxygen species (ROS) burst of leaf discs from 4-week-old WT, *spy-4*, and *sec-2* plants. n = 12. ±S.E.M was shown as error bar. (D) Titer of *Pst* strain DC3000 in 8-day-old WT, *spy-4*, and *sec-2* mutant seedlings at 0 and 3 days after flood inoculation (DAI) of 1 x 10^6^ CFU/mL. Each biological replicate includes 3 seedlings, and 12 biological replicates (n = 12) were used. CFU: Colony forming units. (E) K-means clustering of the 205 core flg22-induced immune genes in 10-d-old WT and *spy-4* seedlings treated with 100 nM flg22 for 1 hour. This gene set represents the intersection of genes upregulated at 10, 30, 90, and 180 minutes post-flg22 treatment, as defined by Bjornson et al.^59^. Expression values are Z-score normalized counts per million (CPM). n = 3 biological replicates. (F) Relative expression level of PTI marker genes *WRKY22* and *WRKY29* in 10-day-old WT, *spy-4*, and *sec-2* mutants treated with 100 nM flg22 for 1 hour (n = 3 biological replicates). **** indicates p < 0.0001, *** indicates p < 0.001, ** indicates p < 0.01, n.s. indicates p > 0.05 with Tukey’s multiple comparisons test.

To determine whether hypersensitivity to elicitors enhances disease resistance in *spy* and *sec* mutants, we inoculated wild-type and mutant seedlings with the phytopathogenic bacterium *Pseudomonas syringae* pathovar *tomato* strain DC3000 (*Pst* DC3000) and measured bacterial growth 3 days post-inoculation. We found that both *spy* and *sec* mutants are more resistant to *Pst* DC3000 compared to wild type, with *spy* mutants showing stronger resistance than the *sec* mutants (Figure 1D; Figure S1C). The phenotypes of *spy-23* and *sec-5* were rescued by transgenes encoding SPY-GFP and SEC-TurboID, respectively (Figure S1C). Taken together, these results indicate that SPY and SEC are required to suppress plant responses associated with MAMP perception and pathogen infection.

To determine how the *spy* mutation alters flagellin-induced gene expression, we performed RNA-seq analysis of wild-type and *spy-4* mutant seedlings after treatment with flg22 for 1 hour. The results demonstrate that *spy-4* is hyper-responsive to flg22 at the global transcriptome level. Among the 205 core flg22-induced genes, 189 (92%) showed a higher average induction in *spy-4* compared to wild-type samples treated with flg22, and 46 (23%) were induced greater than 2-fold in *spy-4* (Figure 1E, Table S2). Quantitative reverse transcription PCR (qRT-PCR) analysis confirmed higher flg22 induction of *WRKY22* and *WRKY29*, two genes encoding pathogen-responsive transcription factors, in *spy-4* compared to wild-type seedlings (Figure 1F). The *sec-2* mutant showed a mild increase in *WRKY29* but similar WRKY22 expression compared to wild type (Figure 1F). These results demonstrate that loss of *O*-glycosylation enhances PTI signaling and disease resistance, with SPY-mediated *O*-fucosylation playing a more prominent role in this process than SEC-mediated *O*-GlcNAcylation.

### SPY and SEC inhibit flg22-induced MAP kinase activation

Flagellin induces PTI through a receptor kinase signaling cascade, from the FLS2 receptor kinase to BIK1 kinase, calcium channels, BSU phosphatase, MAP kinases, ROS burst, and transcription factors.^3,34^ To identify the component of the FLS2 signaling pathway affected by SPY, we tested the effect of SPY inactivation on flg22-triggered downstream signaling events. We found that inhibiting SPY enzyme activity using the chemical inhibitor SOFTI (cite) had no effect on flg22-induced cytosolic calcium increase or phosphorylation of BIK1 (Figure 2A-B), suggesting that SPY may glycosylate components downstream of BIK1. We then analyzed the phosphorylation state of downstream MAP kinases and observed hyperphosphorylation of MAP kinase 3 (MPK3) and MPK6 in flg22-treated *spy-4* and *spy-23* seedlings compared to wild type (Figure 2C-D). The *sec-2* and *sec-5* mutants showed a milder increase in MPK3/6 phosphorylation compared to *spy-4* (Figure 2C, S2A), consistent with the *sec* mutant’s overall weaker phenotypes compared to *spy* (Figure 1). In contrast to the *spy* mutants, a transgenic line that overexpresses SPY-GFP in the *spy-23* background (Figure S2B) showed a weaker MPK3/6 phosphorylation than wild type (Figure 2D, S2A). These findings reveal that SPY is required to inhibit flg22-induced MAP kinase activation, and overexpression of SPY is sufficient to reduce the phosphorylation state of MPK3/6.

**Figure 2.**
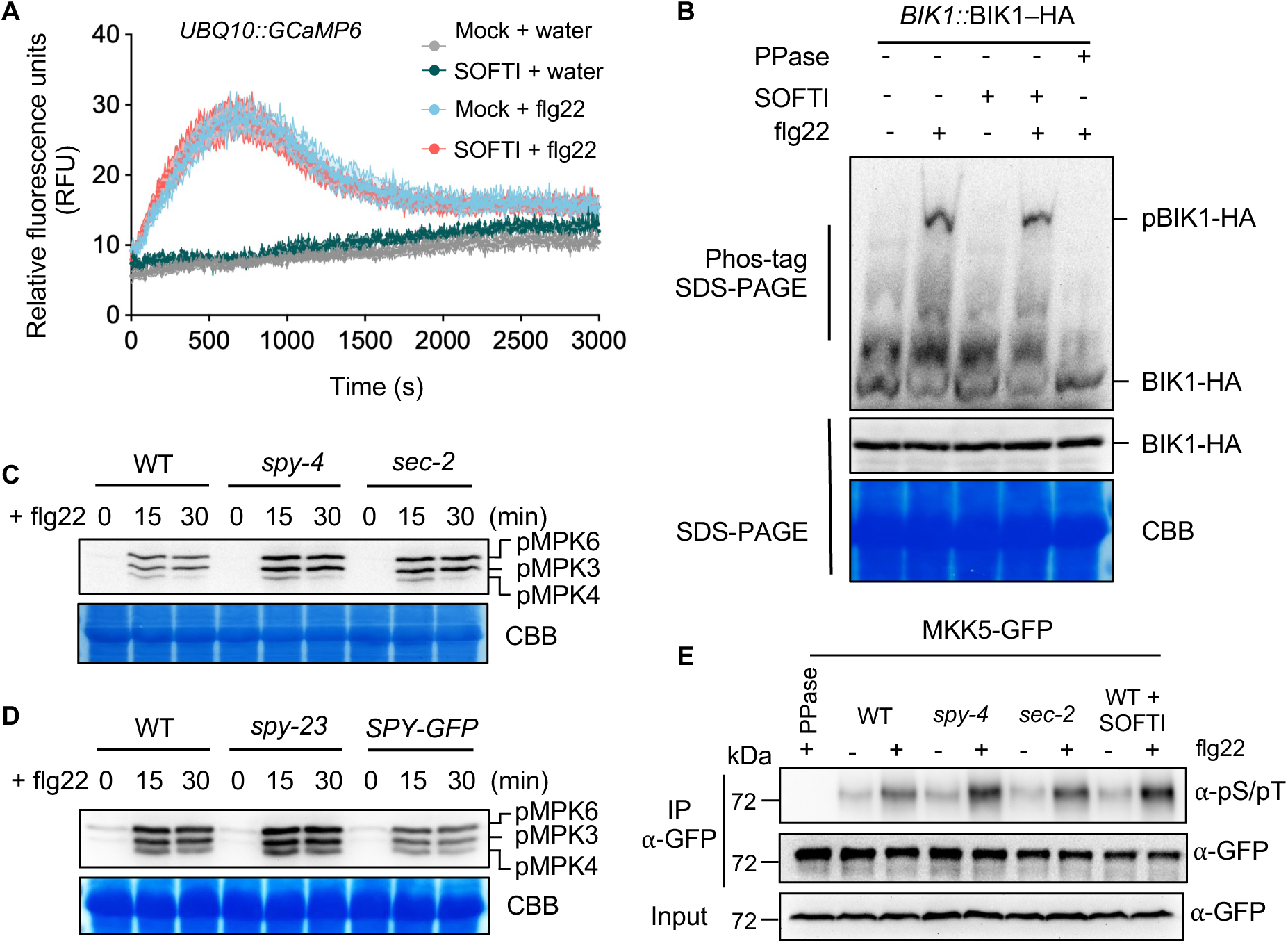
O-glycosylation negatively regulates MAPK cascade activation. (A) Real-time recording of flg22-induced cellular calcium influx measured by GCaMP6 fluorescence calcium sensor in the roots of 10-day-old *UBQ10::GCaMP6* seedlings grown on ½ MS medium supplied with or without 20 μM SPY inhibitor SOFTI. (B) Immunoblots of Phos-tag SDS-PAGE and regular SDS-PAGE showing flg22-induced phosphorylation of BIK1-HA proteins in 10-day-old *BIK1::BIK1-HA* seedlings grown on ½ MS medium supplied with or without 20 μM SOFTI. 1 μM flg22 was applied for 5 minutes. One sample was treated with Lambda phosphatase (PPase) to dephosphorylate BIK1-HA proteins. Immunoblot was probed with anti-HA antibody. (C) Immunoblot showing MAPK phosphorylation level in 10-day-old WT, *spy-4*, and *sec-2* mutant seedlings following treatment with 1 μM flg22 (0-30 min). (D) Immunoblot shows MAPK phosphorylation level in 10-day-old WT, *spy-23*, and *SPY-GFP/spy23* seedlings treated with 1 μM flg22 peptide for the indicated time. Coomassie Brilliant Blue staining (CBB) was used as the loading control. (E) Immunoblot shows flg22-induced phosphorylation of MKK5-GFP in 10-d-old MKK5-GFP/WT, MKK5-GFP/*spy-4*, MKK5-GFP/*sec-2*, and MKK5-GFP seedlings treated with SOFTI. Seedlings were treated with mock or 1 μM flg22 for 5 minutes. MKK5-GFP protein was immunoprecipitated (IP) using anti-GFP magnetic beads and probed with anti-GFP or anti-pS/pT antibody. kDa: kilodalton.

In addition to flagellin, MPK3 and MPK6 are activated by multiple signals, including the fungal elicitor chitin, the bacterial MAMP elf18 (peptide derived from elongation factor Tu), and the endogenous Arabidopsis damage-associated signals RALF1 and pep1 peptides.^35,36^ As with flg22 elicitation (Figure 1A-B), the pep1 peptide inhibited root growth more strongly in *spy-4* and *sec-5* seedlings than in wild type (Figure S3A-B), demonstrating that SPY and SEC are also required to inhibit pep1-mediated responses. Analysis of MPK3/6 activation in wild-type and *spy-4* seedlings treated with pep1 revealed that the phosphorylation state of MPK3/6 was higher in *spy-4* seedlings compared to wild type (Figure S3D). Similar results were observed when additional signals were tested (i.e., chitin, elf18 and RALF1) (Figure S3C-D). These findings reveal that the SPY is required to modulate the outputs (e.g., growth and immunity) of multiple MAMP elicitors.

Next, we tested whether the *spy* and *sec* mutations alter flg22-induced phosphorylation of MAP kinase kinase 5 (MKK5), a direct upstream activator of MPK3/6. To this end, we crossed the MKK5-GFP transgenic line into the *spy-4* and *sec-2* backgrounds to create MKK5-GFP/*spy-4* and MKK5-GFP/*sec-2* lines, respectively. We then treated the respective seedlings with flg22 for 5 minutes, immunoprecipitated MKK5-GFP, and performed immunoblot analysis using an anti-phosphoserine/threonine antibody. Figure 2D shows that flg22 induced higher levels of MKK5 phosphorylation in *spy-4* than in wild-type seedlings. Moreover, inhibition of SPY activity in the MKK5-GFP/WT line using the chemical SOFTI enhanced flg22-induced MKK5 phosphorylation (Figure 2E). These results indicate that SPY-dependent *O*-fucosylation suppresses MKK5 activation during PTI signaling *in vivo*. They also suggest that *O*-fucosylation plays a more prominent role in regulating MKK5 signaling than O-GlcNAcylation.

### SPY directly interacts with and *O*-glycosylates MKK4 and MKK5

Given that previous proteomic studies revealed that MKK4 and MKK5 are modified with fucose^30^, we hypothesized that one or both kinases are SPY substrates. First, we tested whether MKK5 is *O*-fucosylated in wild-type seedlings but not in *spy*-*4* mutant seedlings. We affinity-purified *O*-fucosylated proteins from transgenic plants expressing MKK5-GFP in the wild type and *spy-4* backgrounds using *Aleuria Aurantia* Lectin (AAL), which binds specifically to *O*-fucose. Immunoblot analysis with an anti-GFP antibody showed that AAL enriched MKK5-GFP from wild-type but not in *spy-4* seedlings (Figure 3A). Conversely, MKK5-GFP was immunoprecipitated by an anti-GFP antibody and then analyzed by AAL-biotin gel blotting. Again, we found that *O*-fucosylated MKK5-GFP could be isolated from wild-type but not in *spy-4* seedlings (Figure S4). These results demonstrate that MKK5 is *O*-fucosylated *in vivo* in wild-type but not in the *spy-4* seedlings.

**Figure 3.**
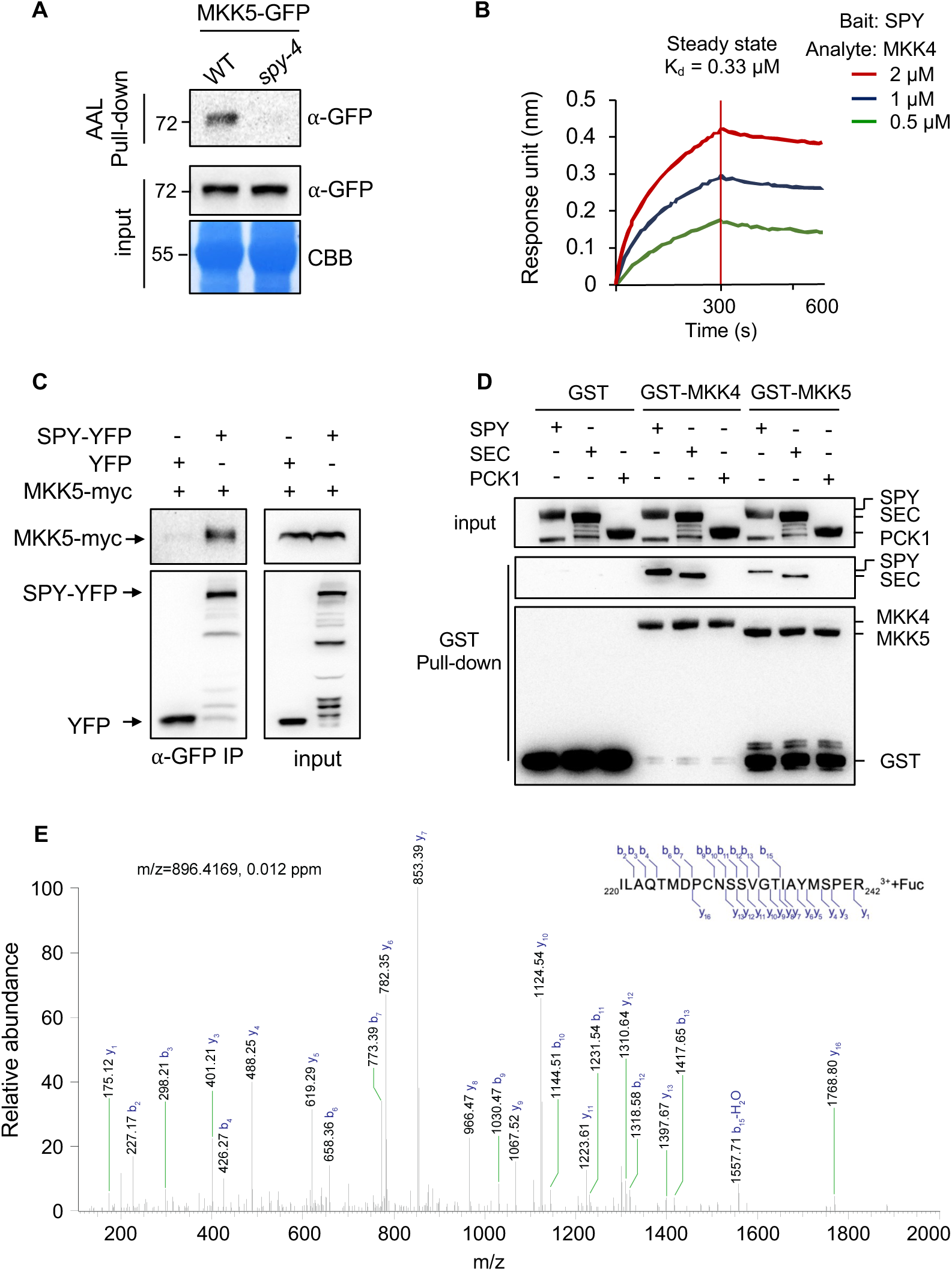
SPY directly interacts with and O-glycosylates MKK4 and MKK5. (A) MKK5-GFP was affinity purified from 7-day-old WT but not *spy-4* seedlings using Aleuria Aurantia Lectin (AAL) immobilized on agarose beads. Coomassie Brilliant Blue staining (CBB) of the membrane shows the loading control. (B) Bio-layer interferometry (BLI) assay showing binding kinetics between SPY and MKK4. 6xHis-SUMO-3TPR-SPY was immobilized to anti-His sensors and dipped into indicated concentrations of GST-MKK4 protein. The dissociation constant (K_d_) was calculated by steady-state fitting of GST-MKK4 concentration against the response unit. (C) Co-immunoprecipitation shows *in vivo* interaction between SPY and MKK5. YFP or SPY-YFP was transiently co-expressed with MKK5-myc in *N. benthamiana* leaves, immunoprecipitated by anti-GFP antibody, and immunoblotted with anti-GFP or anti-myc antibody. (D) *in vitro* affinity purification (pull-down) assay shows direct interaction between SPY/SEC and MKK4/5. Recombinant GST, GST-MKK4, and GST-MKK5 were incubated with 6xHis-tagged SPY, SEC, and PCK1, affinity-purified by Glutathione-agarose beads, and immunoblotted using anti-His and anti-GST antibodies. (E) Higher energy collisional dissociation (HCD) mass spectrum shows O-fucosylation on the activation loop peptide of MKK4 spanning amino acids 220-242, after *in vitro* O-fucosylation by SPY.

We further characterized the direct interactions of SPY with MKK4 and MKK5 using biolayer interferometry (BLI) assays. SPY exhibited a stronger interaction with MKK4 (dissociation constant K_d_ = 330 nM) and a weaker interaction with MKK5 (K_d_ = 560 nM) (Figures 3B and S5A). Co-immunoprecipitation assays confirmed that SPY interacts with MKK5 *in vivo* (Figure 3C). We then analyzed SPY and SEC interactions with MKK4 and MKK5 using *in vitro* pulldown assays. Both GST-tagged MKK4 and MKK5 immobilized on glutathione agarose beads pulled down His-tagged SPY and SEC but not phosphoenolpyruvate carboxykinase 1 (PCK1), a negative control (Figure 3D). These data indicate that SPY and SEC interact with MKK4 and MKK5. After incubation with SPY and GDP-fucose, GST-MKK5 was pulled down by AAL-agarose beads. Omitting GDP-fucose abolished GST-MKK5 binding to AAL, and addition of the SPY inhibitor SOFTI abolished GST-MKK5 binding to AA (Figure S5B). These *in vitro* experiments demonstrate that SPY directly interacts with and O-fucosylates MKK5.

To characterize the *O*-glycosylation sites in MKK4 and MKK5, we performed mass spectrometry analyses of MKK4 and MKK5 after *in vitro O-*glycosylation by SPY and SEC. A conserved peptide, ILAQTMDPCNSSVGTIAYMSPER, in both MKK4 and MKK5 was found to be *O*-fucosylated by SPY and *O*-GlcNAcylated by SEC (Figure 3E, S6). Taken together, these results show that SPY and SEC directly interact with and *O*-glycosylate a conserved peptide in MKK4 and MKK5.

### *O*-glycosylation of residues in activation loop of MKK4 and MKK5 blocks their phosphorylation and activation

As the *O*-fucosylated and *O*-GlcNAcylated peptides in MKK4 and MKK5 contain a S/T-X_3-5_-S/T motif known to be phosphorylated by upstream MEKKs/MAPKKKs for activation,^8,36,37^ we hypothesized that *O*-glycosylation of residues in the activation loop of MKK4 and MKK5 might directly prevent with their phosphorylation or hinder phosphorylation at adjacent sites. To test this, we glycosylated recombinant GST-MKK5 protein by incubating it with SPY or SEC. After affinity purification of GST-MKK5 to remove SPY and SEC, we incubated GST-MKK5 with a truncated, constitutively active MEKK1 in a protein phosphorylation assay. The results in Figure 4A show that preincubation with SPY or SEC in the presence of their donor substrates, GDP-fucose or UDP-GlcNAc respectively, but not in the absence of the substrates, quantitatively reduced the phosphorylation of GST-MKK5 by MEKK1. These results demonstrate that *O*-glycosylation of the activation loop of MKK5 blocks MEKK1-mediated phosphorylation of MKK5.

**Figure 4.**
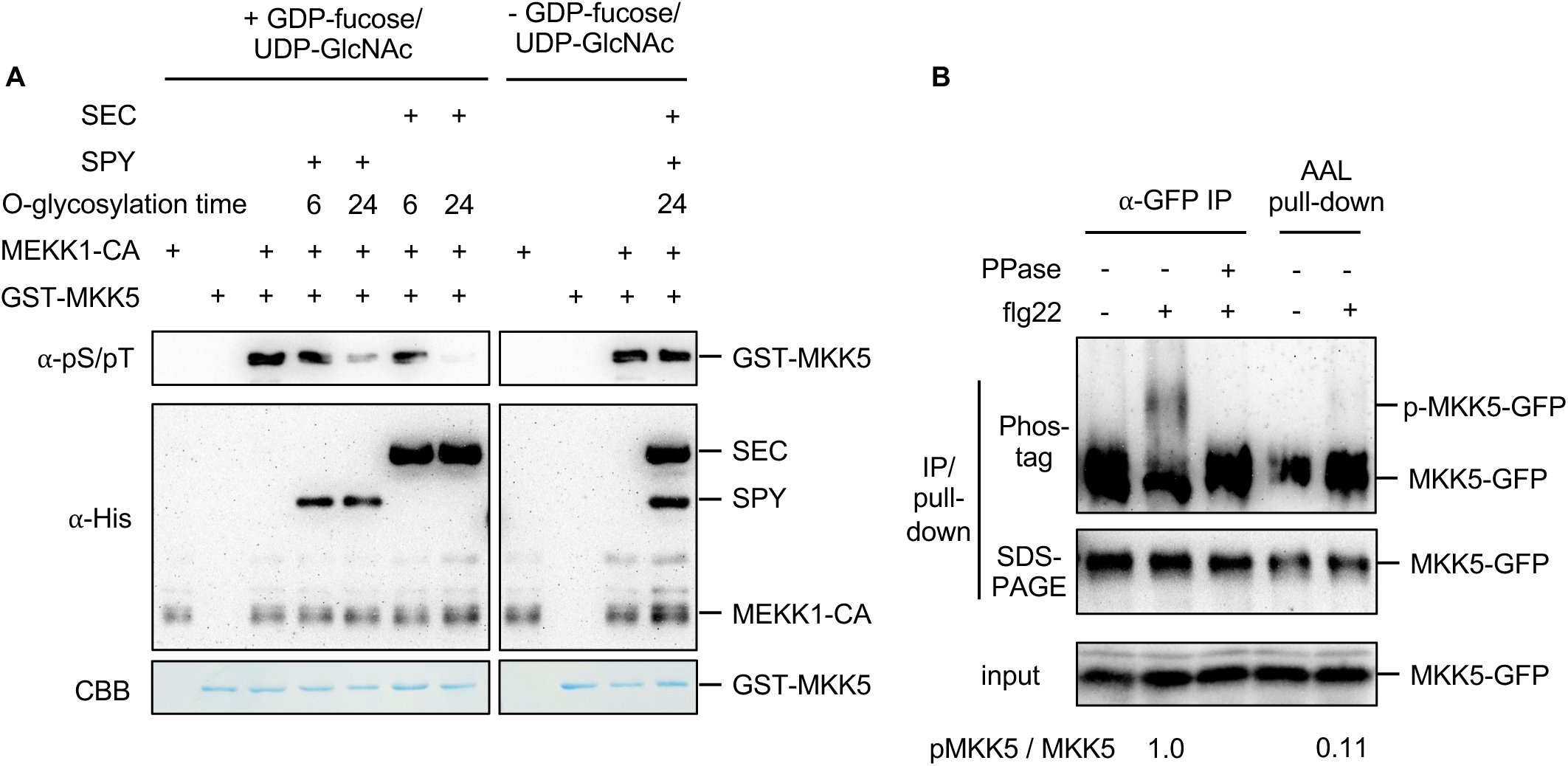
O-glycosylation blocks phosphorylation of MKK5 by MEKK1. (A) Immunoblots show *in vitro* phosphorylation of recombinant GST-MKK5 protein by constitutively active MEKK1-CA kinase before or after O-glycosylation by 6xHis-SUMO-3TPR-SPY and/or 10xHis-MBP-5TPR-SEC for the indicated time with or without the addition of sugar donor GDP-fucose/UDP-GlcNAc. Blots were probed with anti-pS/pT antibody. The gel blot was stained with Coomassie Brilliant Blue (CBB) to show protein loading. (B) AAL pull down preferentially unphosphorylated MKK5. Ten-day-old MKK5-GFP seedlings were treated with mock or 1 μM flg22 for 5 minutes. MKK5-GFP protein was immunoprecipitated using anti-GFP magnetic beads (⍺-GFP IP), and then an aliquot was treated with phosphatase (PPase). An aliquot of the same extract was affinity purified using Aleuria Aurantia Lectin immobilized on agarose beads (AAL pull-down). Immunoblot from Phos-tag gel shows the levels of phosphorylated (p-MKK5-GFP) and unphosphorylated MKK5-GFP.

To test if *O*-fucosylation of MKK5 blocks its phosphorylation *in vivo*, and if flg22 induces phosphorylation of only non-glycosylated MKK5, we treated *MKK5-GFP* seedlings with flg22 to induce phosphorylation of a fraction of MKK5-GFP, which can be detected as a shifted band in the Phos-tag gel blot that is reversed by phosphatase treatment. After AAL pull-down of O-fucosylated MKK5-GFP from the extracts, Phos-tag gel blot showed enrichment of the unphosphorylated MKK5-GFP and depletion of the phosphorylated MKK5-GFP (Figure 4B). These data indicate that phosphorylated MKK5 proteins are not *O*-fucosylated. They also indicate that *O*-fucosylated MKK5 proteins remain unphosphorylated following flg22 elicitation. Together, these findings reveal that *O*-fucosylation of MKK5 blocks its phosphorylation *in vivo*.

### Starvation and pathogen infection reduce protein *O*-glycosylation level

During infection, pathogens deplete host cells of nutrients, including sugar, through various mechanisms^19,38^. Therefore, we hypothesized that changes in sugar availability in plant tissues during pathogen infection may reduce protein *O*-glycosylation levels, allowing plants to fully activate their immune response. To test this, we first determined if sugar availability affects protein *O*-fucosylation levels. We transferred 10-day-old light-grown seedlings into sugar-free media in the dark for 60 hours to deplete endogenous sugar produced by photosynthesis. We then analyzed *O*-fucosylated proteins by immunoblot analysis using an anti-fucose antibody. The results showed that the dark treatment reduced protein *O*-fucosylation levels (Figure 5A). Supplementing sucrose at 6 and 24 hours before the end of the dark treatment recovered *O*-fucosylation of different proteins to various degrees, indicating protein-specific kinetics of *O*-fucosylation upon sucrose treatment (Figure 5A). These results indicate that *O*-fucosylation is sugar-dependent.

**Figure 5.**
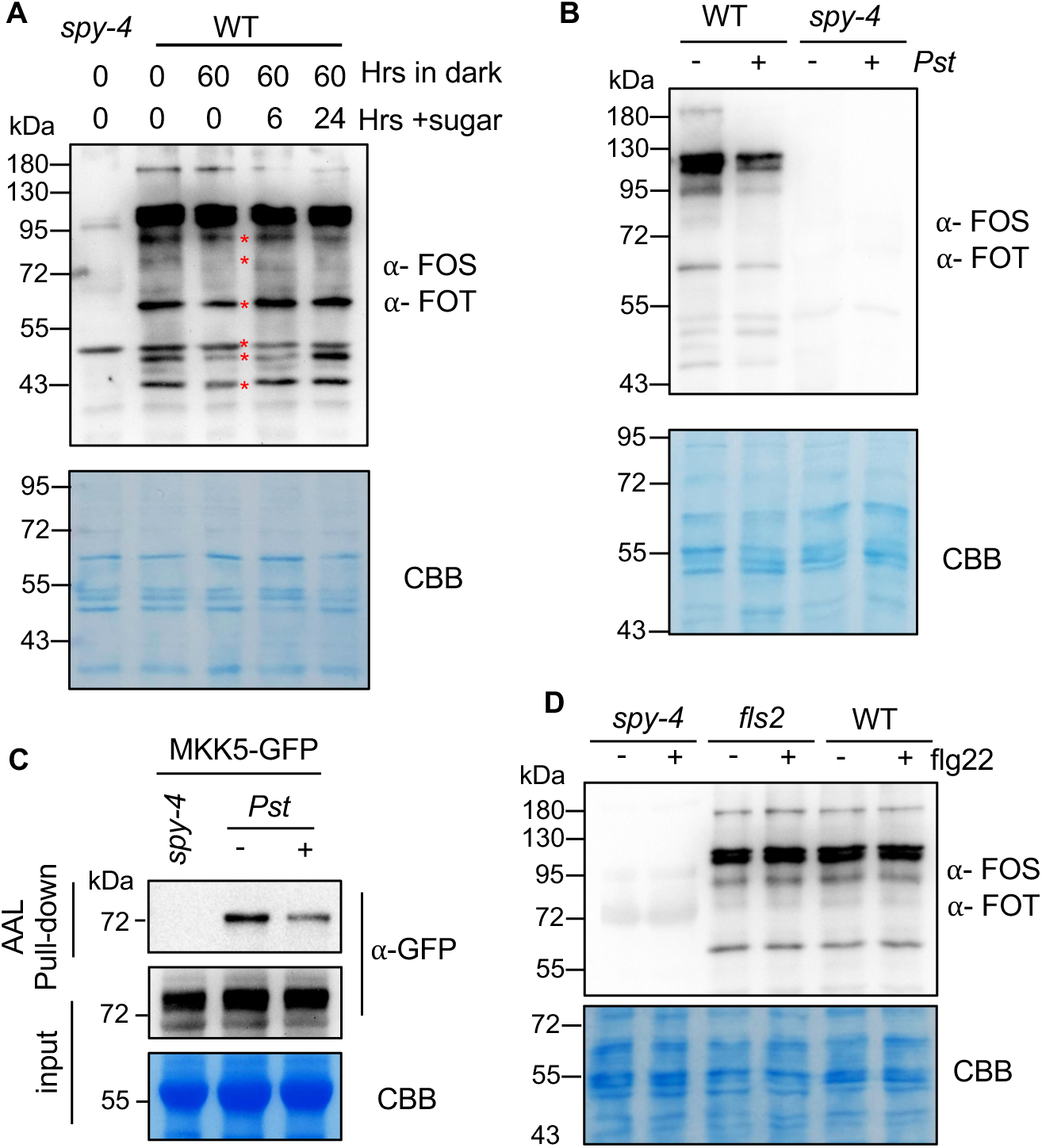
O-fucosylation is decreased by starvation and pathogen infection. A) Immunoblot analysis of O-fucosylated nuclear proteins in WT and *spy-4* seedlings that were grown under light for 10 days and transferred into sugar-free liquid medium in darkness for 60 hours. Sucrose (1%) was added for the indicated time before the end of dark period. The blots were probed with a mixture of anti-Serine-O-fucose (⍺-FOS) and anti-Threonine-O-fucose (⍺-FOT) antibodies and stained with Coomassie Brilliant Blue (CBB). Red asterisks mark the bands with decreased intensity after 60 hours of starvation. (B) WT and *spy-4* plants were grown for 4.5 weeks under short-day conditions, infiltrated with mock solution or a suspension of 1 x 10^5^ C.F.U./mL *Pst* DC3000 (*Pst*) and harvested 3 days post inoculation. Total nuclear protein was immunoblotted with a mixture of anti-Serine-O-fucose (⍺-FOS) and anti-Threonine-O-fucose (⍺-FOT) antibodies. (C) Transgenic plants expressing MKK5-GFP were treated as in panel B. O-fucosylated MKK5-GFP was affinity-purified using AAL-agarose beads and analyzed using anti-GFP antibody. (D) WT, *spy-4*, and *fls2* plants were grown for 4.5 weeks under short-day conditions, infiltrated with mock solution or 100 nM flg22, and total nuclear proteins were immunoblotted using ⍺-FOS and ⍺-FOT 3 days post infiltration. ** indicates p < 0.01, n.s. indicates p > 0.05. p-value are determined by unpaired t-test. The gel blot was stained with Coomassie Brilliant Blue (CBB) to show loading. kDa: kilodalton.

To test how pathogen infection alters host protein *O*-fucosylation, we infected leaves of adult plants with *Pst* DC3000 and analyzed *O*-fucosylation of nuclear proteins. Like sugar starvation, pathogen infection reduced overall *O*-fucosylation levels without significantly affecting SPY protein abundance (Figures 5B, S7). AAL affinity purification of MKK5-GFP revealed that *O*-fucosylation of MKK5-GFP protein was lower in pathogen-infected leaves than in uninfected leaves (Figure 5C). In contrast to pathogen infection, flg22 treatment showed no detectable effects on protein *O*-fucosylation (Figure 5D). These results indicate that pathogen infection decreases the *O*-fucosylation levels of host proteins, including MKK5.

To test whether pathogen infection decreased protein *O*-fucosylation due to reduced sugar availability, we treated infected and uninfected leaves with the sugar donor substrate GDP-fucose. The results show that supplementation with GDP-fucose for 6 hours restored the levels of *O*-fucosylation of proteins, including MKK5, in the pathogen-infected tissues but had no significant effect on protein *O*-fucosylation levels in uninfected tissues (Figures 6A, B). These findings demonstrate that reduced host protein *O*-fucosylation in infected tissues results from decreased availability of sugar substrates.

**Figure 6.**
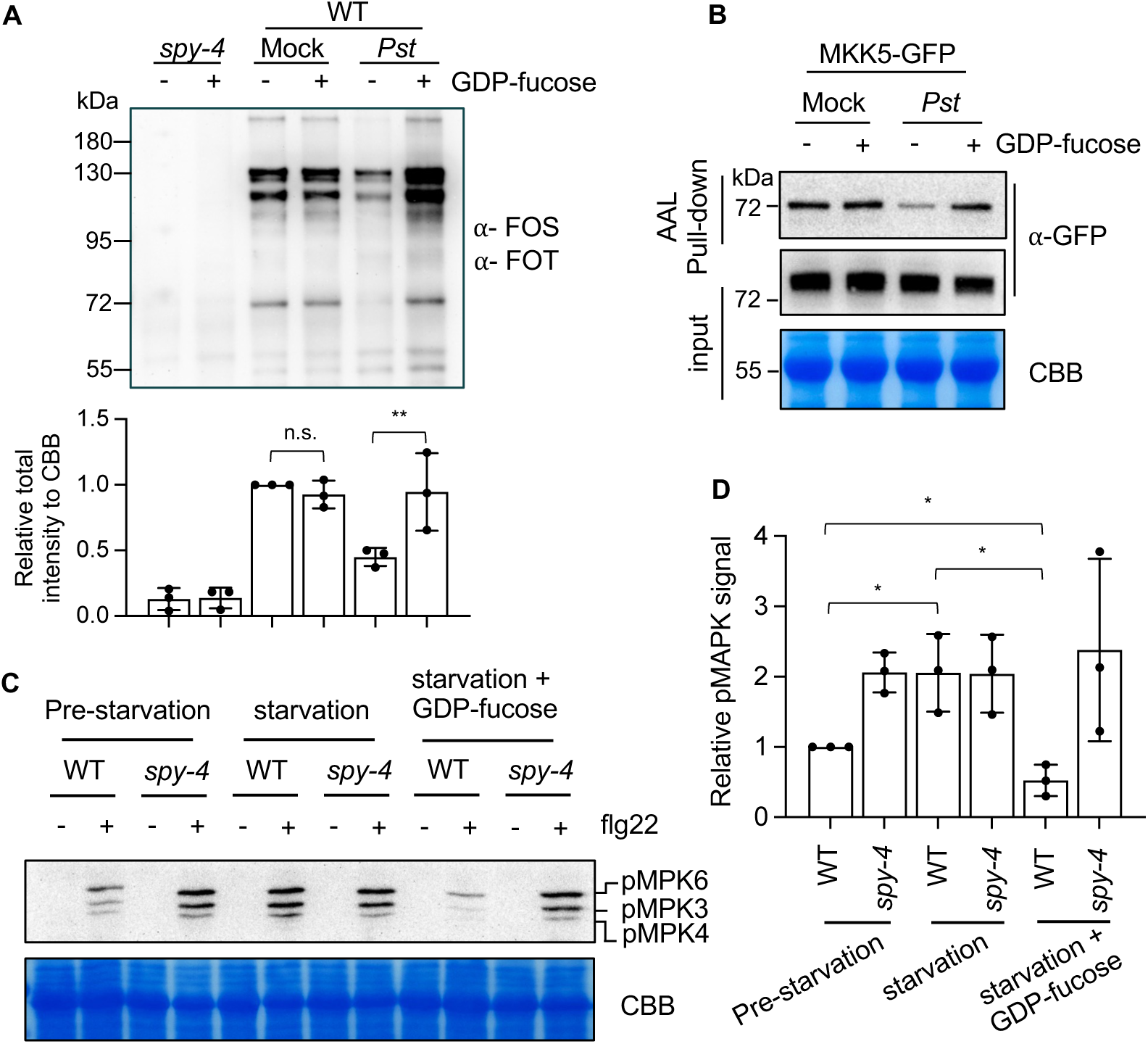
GDP-fucose restores protein O-fucosylation and suppresses flg22-induced MAPK activation in pathogen-infected and starved tissues. (A) Leaves of WT plants were infiltrated with mock solution or a suspension of 1 x 10^5^ C.F.U./mL *Pst*, infiltrated 3 days later with 10 µM GDP-fucose or water, and harvested 6 hours later. O-fucosylated nuclear proteins were immunoblotted using ⍺-FOS and ⍺-FOT. (B) Plants expressing MKK5-GFP were grown and treated as in panel A. O-fucosylated MKK5-GFP was affinity-purified using AAL-agarose beads and analyzed using anti-GFP antibody. (C) WT and *spy-4* seedlings were grown under light for 10 days and transferred into liquid ½ MS medium without sucrose in the dark to cause starvation. GDP-fucose (10 μM) was added after 54 hours, flg22 (1 μM) was added 6 hour later, and seedlings were harvested 15 minutes after flg22 treatment. MPK phosphorylation was analyzed by immunoblotting. (D) Quantitation of relative pMPK signal intensity of flg22-treated samples in (C) normalized to pre-starvation WT plants. n = 3 biological replicates. * indicates p < 0.05. p-value are determined by unpaired t-test. The gel blots were stained with Coomassie Brilliant Blue (CBB) to show loading. kDa: kilodalton.

To test whether SPY-catalyzed O-fucosylation mediates sugar suppression of immune signaling, we analyzed the effects of sugar availability on flg22-induced MPK phosphorylation in wild-type and *spy* plants. Depletion of sugars via 60-hour dark treatment dramatically enhanced flg22-induced MPK3/6 activation in wild-type seedlings, suggesting that starvation enhances immunity. In contrast, starvation had little effect on MPK3/6 in *spy-4* seedlings, which displayed constitutively high levels of MPK3/6 activation (Figures 6C, D, S8A), consistent with starvation enhancing MPK activation by reducing SPY-catalyzed *O*-fucosylation of MKK4/5. Inclusion of 1% sucrose in the media increased MPK activation in both wild type and *spy,* while *spy* remained hyper-responsive to flg22 compared to wild type (Figure S8B). This observation indicates that exogenous sugar represses MPK through a SPY-dependent mechanism but also enhances immunity through a SPY-independent mechanism. Indeed, it was recently reported that exogenous sugar has an immune-enhancing effect via a calcium-dependent kinase.^39^ Consistent with this scenario, GDP-fucose treatment was sufficient to restore MKK *O*-fucosylation (Figure 6A, B) and repress flg22-induced MPK phosphorylation in starvation-treated wild-type plants (Figure 6C, D). Neither starvation nor GDP-fucose treatment had a significant effect on MPK3/6 phosphorylation in the *spy* mutant (Figures 6C-D and S8A). Together, these data demonstrate that sugar availability modulates MKK4/5 *O*-glycosylation and, in turn, MPK3/6 activation through a SPY-dependent mechanism.

### Chemical inhibition of SPY enhances plant defense

To test whether chemical inactivation of SPY would enhance disease resistance, we grew Arabidopsis seedlings on media containing 0 or 20 µM SOFTI and inoculated them with *Pst* DC3000. We found that SOFTI treatment reduced *Pst* DC3000 growth in wild-type seedlings (Figure 7A), whereas SOFTI did not inhibit *Pst* DC3000 growth in liquid culture (Figure S9), indicating that inhibition of SPY-mediated *O*-fucosylation enhances host immunity. In addition, SOFTI treatment showed no significant effect on immunity against *Pst* DC3000 in *spy-4* seedlings, confirming that SOFTI enhances immunity by inhibiting SPY. SOFTI treatment had a significantly weaker effect on *Pst* DC3000 proliferation in the partial loss-of-function *mkk4/5* mutant than in wild type (Figure 7B), consistent with the conclusion that inhibiting SPY activity enhances defense by increasing the level of active MKK4/5. In contrast, SOFTI treatment of the *mkp1* mutant had a normal immune-enhancing effect (Figures 7C and S10). MKP1 (MAP kinase phosphatase 1) dephosphorylates and inactivates MPK3/6, thereby inhibiting immune responses.^40^ Collectively, these results show that chemical inhibition of SPY enhances Arabidopsis disease resistance to *Pst* DC3000 largely by activating MKK4/5.

**Figure 7.**
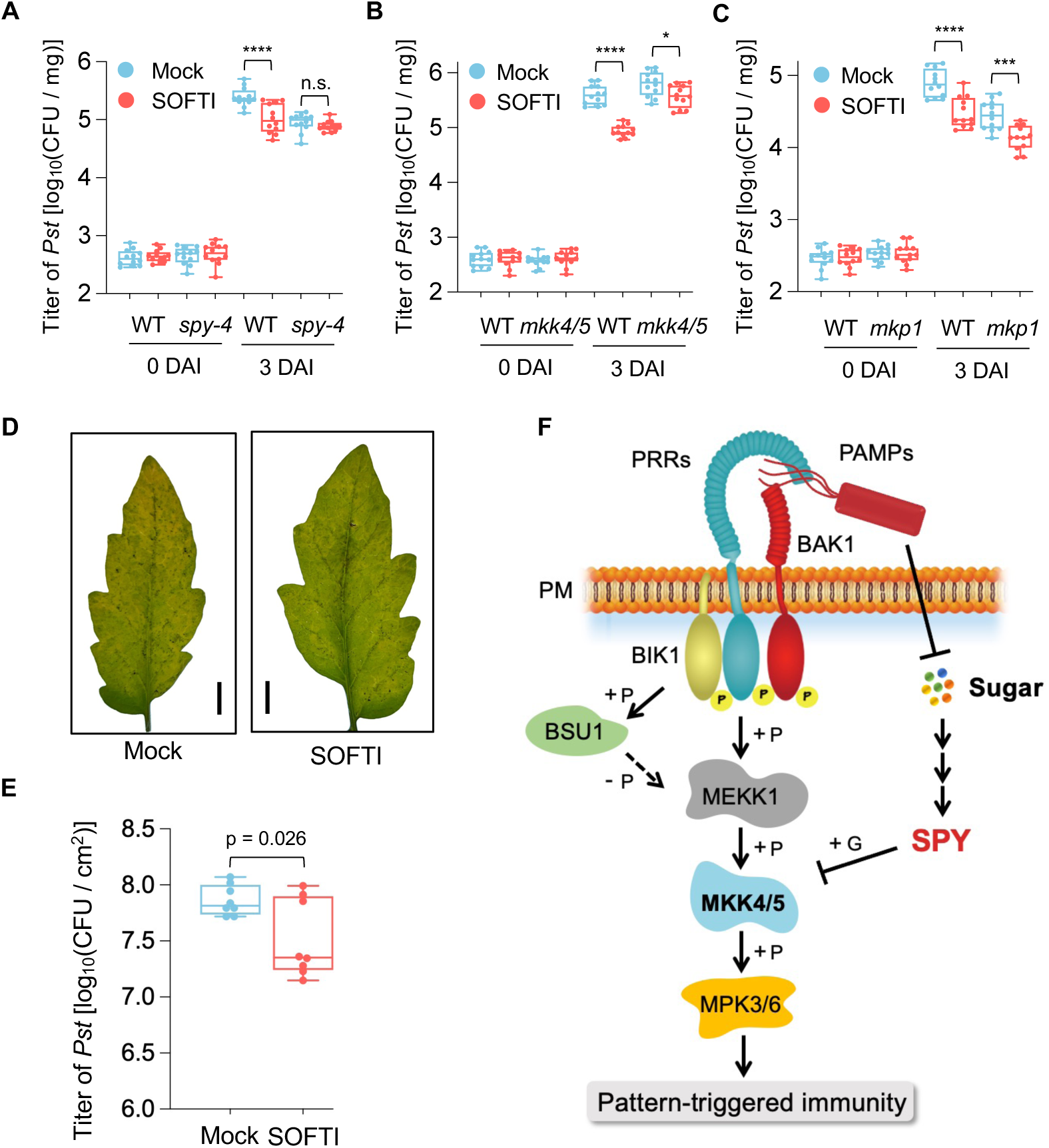
SOFTI inhibition of SPY enhances disease resistance in Arabidopsis and tomato. (A) – (C) Titer of *Pst* DC3000 in flood-inoculated (1×10^6^ CFU / mL) 7-day-old wile-type (WT), *spy-4*, *mkk4-18 mkk5-18* (*mkk4/5*), or *mkp1* seedlings grown on ½ MS medium supplemented with or without 20 µM SPY inhibitor SOFTI, measured at 0 (right after inoculation) or 3 days after inoculation (DAI). Each box plot shows the minimum, first quartile, median, third quartile, and maximum of the colony-forming units per milligram tissue (CFU/mg), from 12 biological repeats, each including three seedlings. **** indicates p < 0.0001, *** indicates p < 0.001, * indicates p < 0.05, n.s. indicates p > 0.05 with Tukey’s multiple comparisons test. The numbers above the asterisk indicate the fold difference between the average CFU/mg of the two groups. (D) Disease symptoms of a pair of tomato leaflets treated with mock or 20 µM SOFTI and then infiltrated with 2 x 10^4^ CFU/mL *Pst* DC3000. Images are representative of n = 5 biological replicates. Bars = 1 cm. (E) Titer of *Pst* DC3000 in tomato leaves infiltrated with 1 x 10^5^ CFU/mL *Pst* DC3000 with or without SOFTI, at 4 DAI. Data are from 2 pairs of leaflets per plant and 4 individual plants with Tukey’s multiple comparisons test. (F) A diagram of the proposed molecular mechanism. Perception of pathogen-derived PAMPs by PRRs (*e.g*. FLS2) triggers phospho-relay leading to immune activation. O-glycosylation of MKK4/5 blocks the phospho-relay and restricts immune response. Pathogen-induced sugar depletion reduces O-glycosylation of MKK4/5, allowing for MKK4/5 phosphorylation and activation, and consequently enhancing the immune response. +P: phosphorylation. -P: dephosphorylation. +G: O-glycosylation. PM: plasma membrane.

We also tested the effect of SOFTI treatment on *Pst* DC3000 infection of susceptible tomato (*S. lycopersicum*) plants. Consistent with the observation in Arabidopsis, SOFTI treatment alleviated disease symptoms and reduced *Pst* DC3000 proliferation in tomato leaves (Figure 7D, E). Taken together, these data show that chemical inhibition of SPY enhances immunity to *Pst* DC3000 in Arabidopsis and tomato plants, providing further evidence for the key role of SPY and protein *O*-glycosylation in the metabolic regulation of immunity (Figure 7F).

## Discussion

Extensive studies of the plant immune system have yielded a deep understanding of the signaling pathways linking microbial perception to resistance responses, including gene expression and metabolic reprogramming.^4^ It has also emerged as a prominent feature of plant pathogens that successful infection often requires the exploitation of nutrients from host cells,^15^ likely due to the absence of circulating body fluids in plants, unlike animals. On the other hand, numerous beneficial microbes enhance nutrient acquisition and are crucial for plant health in nature and agriculture.^1,2^ The microbial impact on host nutrient status appears to be a defining factor in plant-microbe relationships in ecological settings and across different stages of pathogen infection; thus, linking nutrient sensing to immune signaling should be of paramount importance. Our study identifies *O*-glycosylation of MKKs as such a critical link that suppresses immunity under nutrient sufficiency but enhances immunity upon nutrient loss.

We discover that the *spy* and *sec* mutants are hypersensitive to MAMP-induced growth inhibition, gene expression, ROS bursts, and disease resistance, all downstream of MAPKs,^41^ but show normal MAMP-induced calcium burst and BIK1 phosphorylation, which are upstream of MAPKs in the PTI pathways.^34^ The genetic evidence suggests that *O*-glycosylation inhibits the MAPK module. We then demonstrate that SPY and SEC catalyze *O*-fucosylation and *O*-GlcNAcylation, respectively, of the activation loops of MKK4 and MKK5, thereby preventing their phosphorylation and activation by upstream MEKKs/MAPKKKs in the PTI signaling pathway. We further show that starvation reduces MKK4/5 *O-*fucosylation and increases MAMP-induced MPK3/6 phosphorylation; such effect of starvation on MPK3/6 activation is reversed by GDP-fucose treatment, demonstrating that sugar depletion reduces cellular GDP-fucose availability and *O*-fucosylation level of MKK4/5, leading to enhanced immune signaling. Similarly, pathogen infection decreases *O*-fucosylation of MKK4/5, and this effect is reversed by exogenous GDP-fucose, confirming that reduced availability of the sugar substrate leads to decreased O-fucosylation of MKK4/5 in infected plant cells. Pathogen infection may reduce host GDP-fucose levels by exploiting sugar precursors or altering metabolic pathways involved in GDP-fucose synthesis. Corroborating the genetic and biochemical evidence for the immune-suppressing function of *O*-glycosylation, chemical inhibition of SPY enhances MAMP-induced MAPK activation and resistance to a bacterial pathogen.

*O*-glycosylation may suppress immunity by modulating additional immune signaling components. In addition to MKK4/5, proteomic profiling has revealed *O*-glycosylation of several other immunity-related proteins, including MAP kinase phosphatase 1 (MKP1).^30^ MKP1 dephosphorylates MKP3/6 and negatively regulates immunity, and the *mkp1* mutant shows increased resistance to *Pst* DC3000 infection.^40^ *O*-glycosylation may potentially activate MKP1 to suppress immune responses. However, unlike the *mkk4/5* mutant, which exhibited a reduced immune response to SOFTI treatment, *mkp1* showed a similar SOFTI-enhanced pathogen resistance to the wild type (Figure 7C and S10), suggesting that MKP1 is not essential for SPY *O*-fucosylation to modulate the defense response. However, these data do not rule out a minor contribution from MKP1 *O*-glycosylation, in parallel with MKK4/5. SOFTI treatment also had no noticeable effect on flg22-induced BIK1 phosphorylation and cytosolic calcium increase, which is mediated by BIK1 phosphorylation of the calcium channels CNGC2/4,^34^ suggesting that SPY *O*-fucosylation does not affect upstream flg22/FLS2 signaling. Together, these results suggest that MKK4/5 are the major targets for SPY regulation of PTI signaling.

Collectively, our findings support the model of metabolic regulation of immunity: *O*-glycosylation of MKKs acts as a metabolic rheostat that fine-tunes immune responses according to sugar availability during plant-microbe interactions. Under normal conditions, portions of cellular MKK4 and MKK5 are *O*-glycosylated and cannot be activated, thereby restraining PTI. Reduced GDP-fucose availability during pathogen infection decreases MKK4/5 *O*-glycosylation, thereby increasing MKK4/5 phosphorylation and activity, resulting in enhanced immune responses (Figure 7F).

The link between sugar sensing and immune signaling is likely to play an important role in optimizing the spatiotemporal dynamics of immune responses during the progression of infection. As infection initiates locally, host cells in contact with and exploited by pathogens would need to mount a more intense defense response, such as cell death, than those in the vicinity do. Such spatial heterogeneity of immune states has been revealed by recent single-cell transcriptomic analyses of Arabidopsis leaves infected by *Pseudomonas syringae.*^42,43^ However, the mechanisms underlying this immune-state heterogeneity remain unknown. Our study suggests that heterogeneity in pathogen nutrient exploitation may contribute to differences in PTI signaling intensity and immune cell states in infected plant tissues.

Metabolic regulation of PTI signaling intensity may play a broad role in plant interactions with diverse microbes, ranging from beneficial to pathogenic. As MAMPs are conserved across pathogenic and non-pathogenic microbes, PTI signaling must be modulated by other mechanisms to establish appropriate interactions with diverse microbes, namely, by enhancing defense responses against virulent pathogens but dampening PTI signaling during interactions with neutral and beneficial microbes.^1,2^ The mechanism uncovered in our study would contribute to immune suppression under nutrient-sufficient conditions, such as interaction with beneficial microbes, while enhancing immune activation upon nutrient depletion, such as during pathogen infection. Future research should investigate whether O-glycosylation contributes to immune suppression during plant interactions with beneficial microbes that improve host nutrient status.

*O*-glycosylation, catalyzed by OGTs and their homologs, is an essential cellular signaling mechanism in both plants and mammals.^28^ Extensive studies over four decades have shown the broad implications of *O*-GlcNAcylation in human diseases, including diabetes, cancer, neuron degradation, and immunological disorders.^44^ There is a growing body of literature suggesting a key role for *O*-GlcNAcylation in modulating immune responses during infections in mammals, with both proinflammatory and anti-inflammatory effects of *O*-GlcNAc observed across different immune cell types.^24^ Notably, *O*-GlcNAcylation of receptor-interacting kinase 3 (RIPK3), which promotes necroptosis downstream of the leucine-rich repeat Toll-like receptors, inhibits RIPK3 function and suppresses immune activation and necroptosis, analogous to *O*-glycosylation of MKK4/5 inhibiting immunity in plants.^45,46^ In contrast to MKK4/5, which are inactivated by *O*-glycosylation of the kinase activation loop, RIPK3 is *O*-GlcNAcylated in its regulatory domain that mediates protein interaction.^45^ Thus, *O*-glycosylation appears to mediate metabolic regulation of innate immunity in both plants and animals through distinct mechanisms that modulate immune kinases. Furthermore, these observations are likely the tip of the iceberg, as proteomic studies have revealed significant overlaps between *O*-glycosylation and phosphorylation in animal immune cells and in plant cells.^28,47–49^

The identification of *O*-glycosylation as a regulator of immune signaling suggests new strategies to improve plant disease resistance. As SPY and SEC are essential for viability and play key roles in broad developmental processes, genetic alterations in SPY and SEC are likely to have pleiotropic effects. However, temporary chemical inhibition of SPY may achieve beneficial outcomes in fighting pathogen infection. It’s encouraging that SOFTI enhances immunity and reduces pathogen infection in both Arabidopsis and tomato. Future discoveries of high-affinity, low-cost, substrate-specific inhibitors that block SPY-MKK interaction may lead to the development of immune-enhancing agrichemicals. In addition to chemical inhibitors, it is conceivable that improved disease resistance can be achieved by altering MKK *O*-glycosylation through genetic engineering. Our discovery of the molecular link between sugar sensing and immunity signaling suggests new strategies for managing plant-microbe interactions in agriculture.

### Limitations of the study

*O*-glyosylation blocking phosphorylation on MKK5 activation loop was demonstrated *in vitro* and supported by several *in vivo* experiments, including (1) mutual exclusion between *O*-fucosylation and phosphorylation of MKK5 (Figure 4B), and (2) significant loss of SOFTI effect in the *mkk4/5* double mutant, confirming MKK4/5 being the major functional targets of *O*-fucosylation (Figure 7B). However, the causal relationship between *O*-glycosylation and blocking phosphorylation was not demonstrated *in vivo* using traditional site-directed mutagenesis of the modification sites. However, such a mutagenesis experiment is not possible in this case because mutations would block both *O*-glycosylation and phosphorylation. Further, the S/T residues in the activation loop are required for the kinase activity of MKK4/5,^36,37^ and MKK4/5 activity is required for viability.^50^ Future development of tools for substrate-specific manipulation of *O*-glycosylation, such as inducible interactions between MKK4/5 and SPY or an *O*-fucosidase, will enable direct *in vivo* analysis of the effects of altering *O*-glycosylation on MKK4/5 phosphorylation. Future studies will determine how *O*-glycosylation of MKK4/5 optimizes immune responses during plant interactions with diverse microbes, including beneficial symbionts.

## Acknowledgements

We thank Dr. Shuqun Zhang from the University of Missouri for kindly providing the MKK5-GFP transgenic line, Dr. Scott Peck for providing the *mkp1* mutant, and Danbi Byun for generating the *SEC-TbID/sec-5* transgenic plants. This work is supported by the National Institute of General Medical Sciences (NIGMS) under Award Number R01GM066258 to Z.W. and the Carnegie Endowment Fund to the Carnegie Mass Spectrometry Facility.

## Author Contributions

YA, YB, HZ, MBM, and Z-YW designed the project; MBM and Z-YW supervised the project; YA, YB, J-GK performed the pathogen growth assay and recorded the disease phenotype; YA, YB, and AC performed the seedling growth assays; YA, HZ, and YB performed the *in vitro* and *in vivo* biochemistry assays; YA and HZ prepared samples for mass spectrometry, CST, YB, and S-LX analyzed the mass data; YA prepared samples for RNA-sequencing, and QJS analyzed the data; YA and Z-YW drafted the manuscript with input from all authors. MBM and J-GK reviewed and edited the manuscript.

## Data and Materials Availability

All data presented in this study are available from the corresponding author on request.

## Supplemental materials

**Figure S1.**
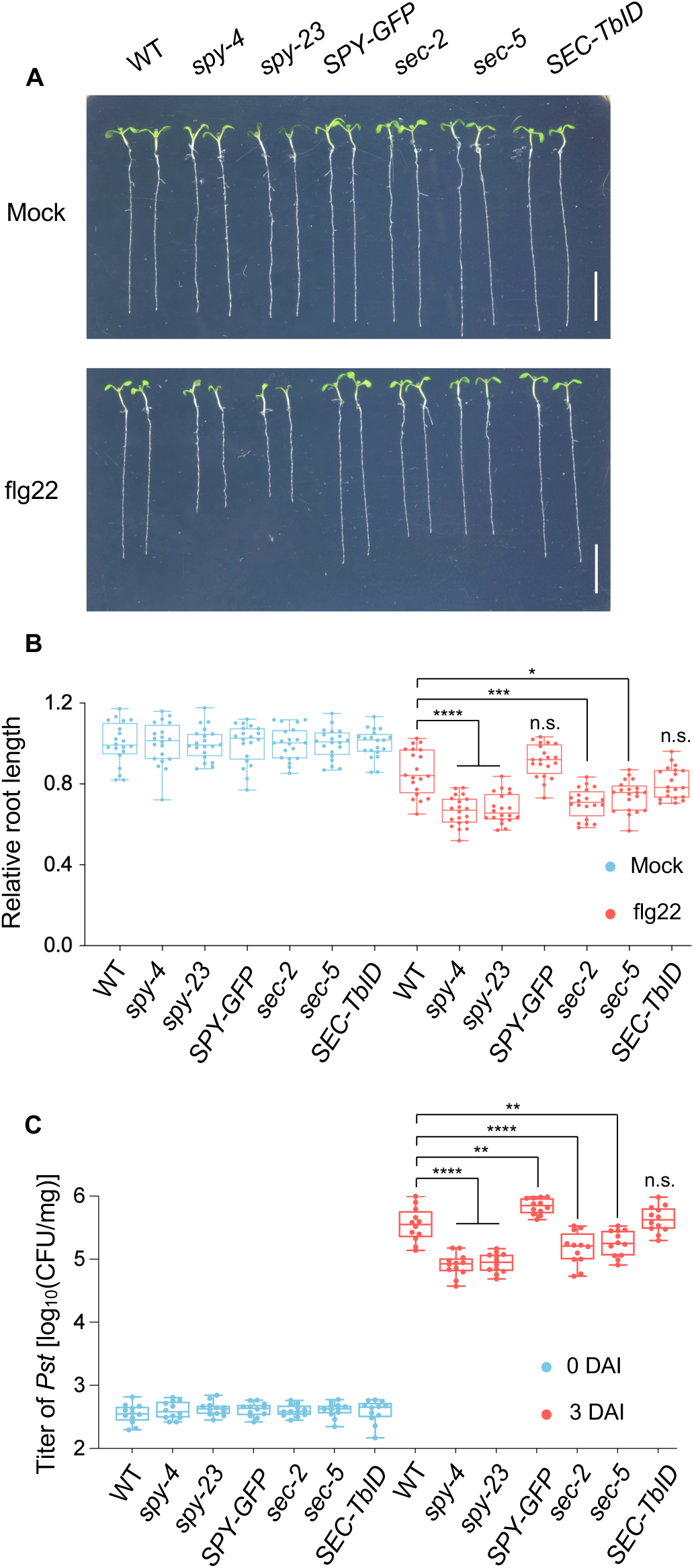
Genetic analysis of SPY and SEC in PTI response. (A) Primary root length phenotype of 9-day-old wild-type (WT), *spy-4*, *spy-23, SPY-GFP/spy-23, sec-2, sec-5,* and *SEC-TbID/sec-5* seedlings grown on ½ MS medium supplemented with or without 1 μM flg22 in short-day conditions. Bar = 1 cm. (B) Quantification of root length of seedlings (n = 20 biological replicates) shown in (A). (C) Titer of *Pst* strain DC3000 in 7-day-old wild-type Col-0, *spy-4*, *spy-23, SPY-GFP/spy-23, sec-2, sec-5,* and *SEC-TbID/sec-5* seedlings at 0 and 3 days after flood inoculation (DAI) of 1 x 10^6^ CFU/mL. Each biological replicate includes 3 seedlings, and 12 biological replicates (n = 12) were used. CFU: Colony-forming units. **** indicates p < 0.0001, *** indicates p < 0.001, ** indicates p < 0.01, * < indicates p < 0.05, n.s. indicates p > 0.05 with Tukey’s multiple comparisons test.

**Figure S2.**
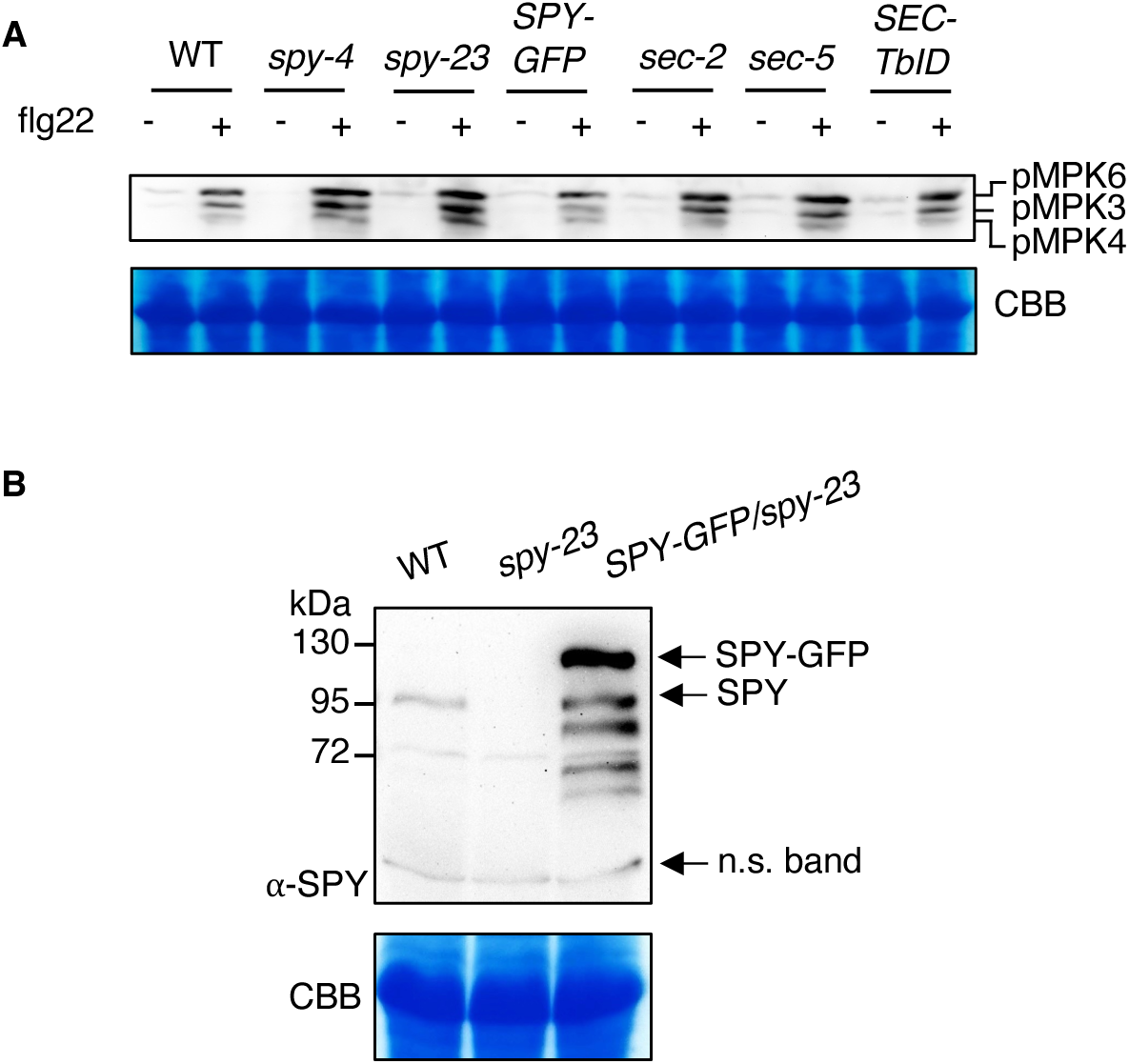
O-glycosylation negatively regulates MAPK cascade activation. (A) Immunoblot showing MAPK phosphorylation in 10-day-old WT, *spy-4*, *spy-23, SPY-GFP/spy-23, sec-2, sec-5,* and *SEC-TbID/sec-5* seedlings treated with 1 μM flg22 peptide for 15 minutes. (B) SPY-GFP is overexpressed in the *SPY-GFP/spy-23* transgenic line. Immunoblot showing the level of SPY proteins in 10-d-old WT, *spy-23*, *SPY-GFP/spy-23* seedlings detected with anti-SPY antibody. kDa: kilodalton. Gel blots were stained with Coomassie Brilliant Blue (CBB) to show protein loading.

**Figure S3.**
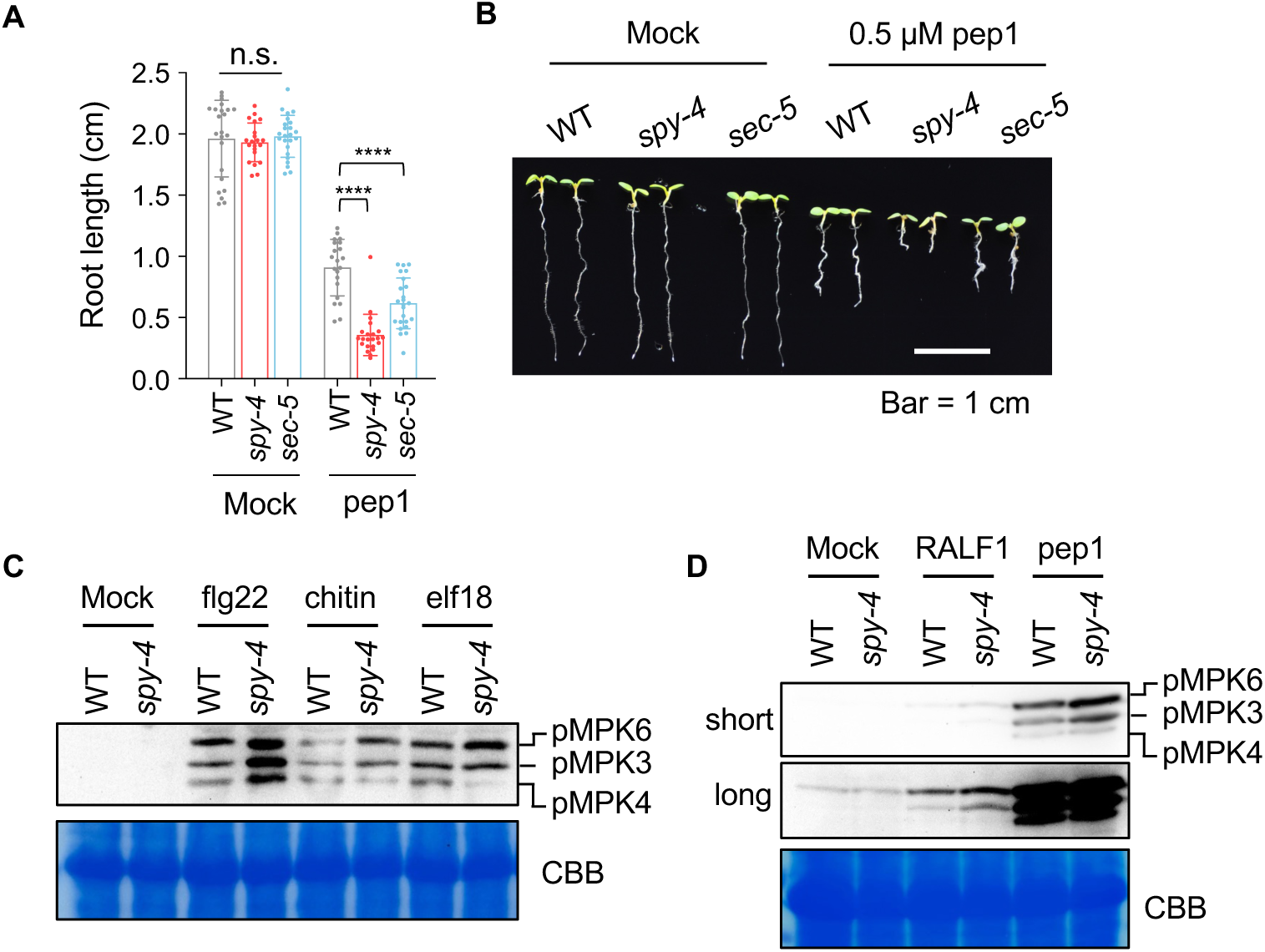
Loss of O-glycosylation increases plant sensitivity to multiple elicitors. (A) Quantification of primary root length of 6-day-old WT, *spy-4*, and *sec-5* mutants grown on ½MS medium supplemented with or without 0.5 μM pep1. **** indicates p < 0.0001, n.s. indicates p > 0.05 with Tukey’s multiple comparisons test. (B) Representative seedlings from treatments shown in (A). (C) and (D) Immunoblot showing MAPK phosphorylation level in 10-day-old WT and *spy-4* seedlings treated by 1 μM flg22, 500 μg/mL chitin, 1 μM elf18, 1 μM RALF1, or 1 μM pep1 for 15 minutes. Short and long exposure are shown. Gel blots were stained with Coomassie Brilliant Blue (CBB) to show protein loading.

**Figure S4.**
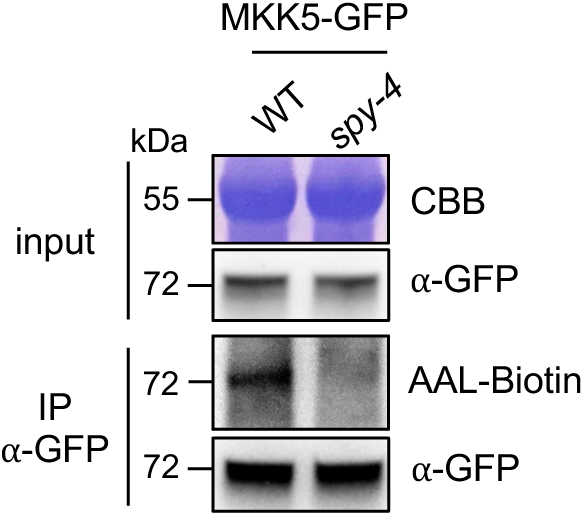
SPY O-fucosylates MKK5 *in vivo*. MKK5-GFP was Immunoprecipitated using anti-GFP beads from protein extracts of 12-day-old MKK5-GFP/WT and MKK5-GFP/*spy-4* seedlings. The gel blots were probed with AAL-biotin and anti-GFP antibody and stained with Coomassie Brilliant Blue (CBB). kDa: kilodalton.

**Figure S5.**
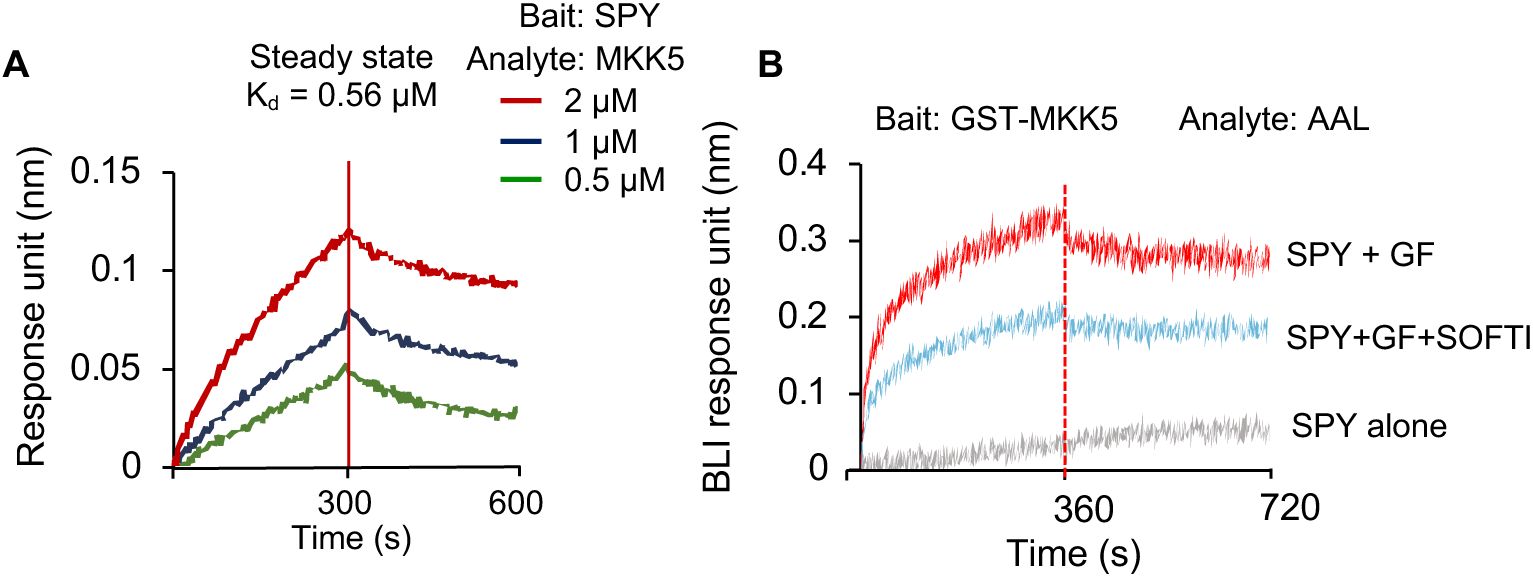
SPY interacts with and O-fucosylates MKK5 *in vitro*. (A) Bio-layer interferometry (BLI) showing binding kinetics between SPY and MKK5. 6xHis-SUMO-3TPR-SPY was immobilized to anti-His sensors and dipped into indicated concentrations of GST-MKK5 proteins. The dissociation constant (K_d_) was calculated by steady-state fitting of GST-MKK5 concentration against the response unit. (B) BLI assay showing interaction between O-fucosylated GST-MKK5 protein with AAL. GST-MKK5 was incubated with SPY *in vitro,* with or without GDP-fucose (GF) and SOFTI, loaded onto anti-GST biosensors, and analyzed for binding to AAL.

**Figure S6.**
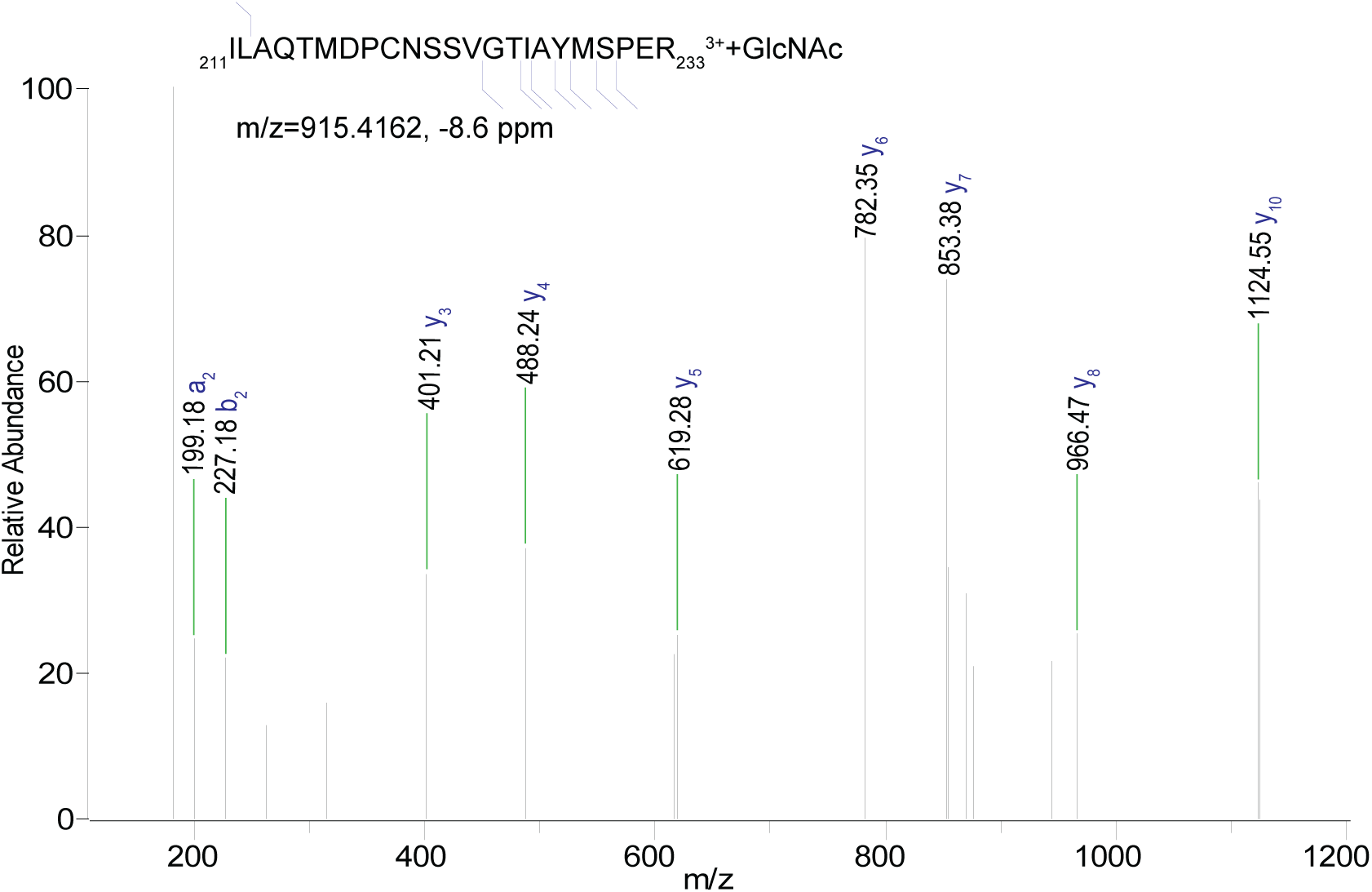
SEC O-GlcNAcylates MKK5 in its kinase activation loop. Higher energy collisional dissociation (HCD) mass spectrum shows O-GlcNAcylation on the activation loop peptide of MKK5 spanning amino acids 211-233, after *in vitro* O-GlcNAcylation by SEC.

**Figure S7.**
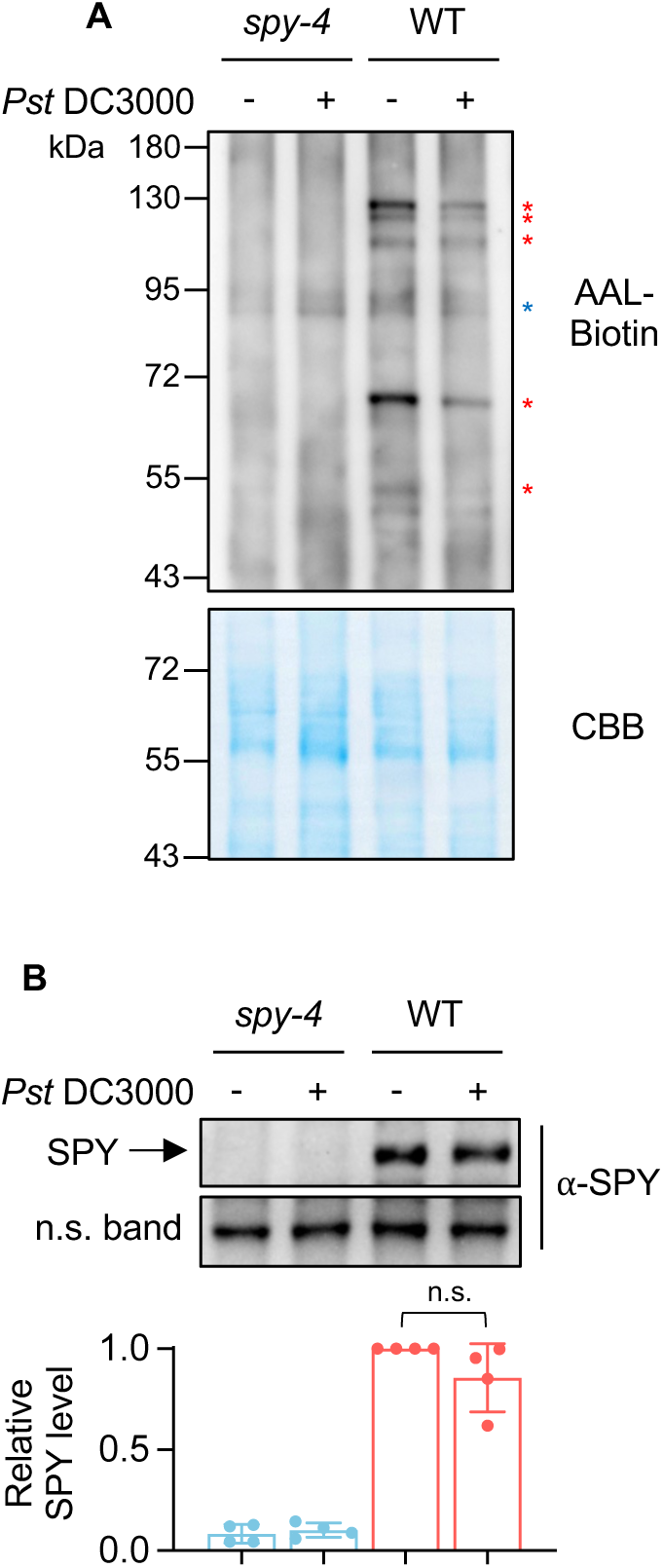
Pathogen infection reduces host protein O-fucosylation. (A) Immunoblot analysis of O-fucosylated nuclear proteins of 4.5-week-old short-day-grown WT and *spy-4* seedlings 3 days after infiltration with mock solution or *Pst* DC3000 (suspension of 1 x 10^5^ CFU/mL). The blot was probed with AAL-biotin and Streptavidin-HRP. Bands of O-fucosylated proteins are marked by red asterisks, and a non-specific band by a blue asterisk. The gel blot was stained with Coomassie Brilliant Blue (CBB) to show loading. kDa: kilodalton. (B) Immunoblot and band intensity quantification (normalized to the non-specific [n.s.] band) showing the level of SPY proteins in the same samples described in (A). n = 4 biological replicates. The blot was probed with anti-SPY antibody. n.s. indicates p > 0.05 with Tukey’s multiple comparisons test.

**Figure S8.**
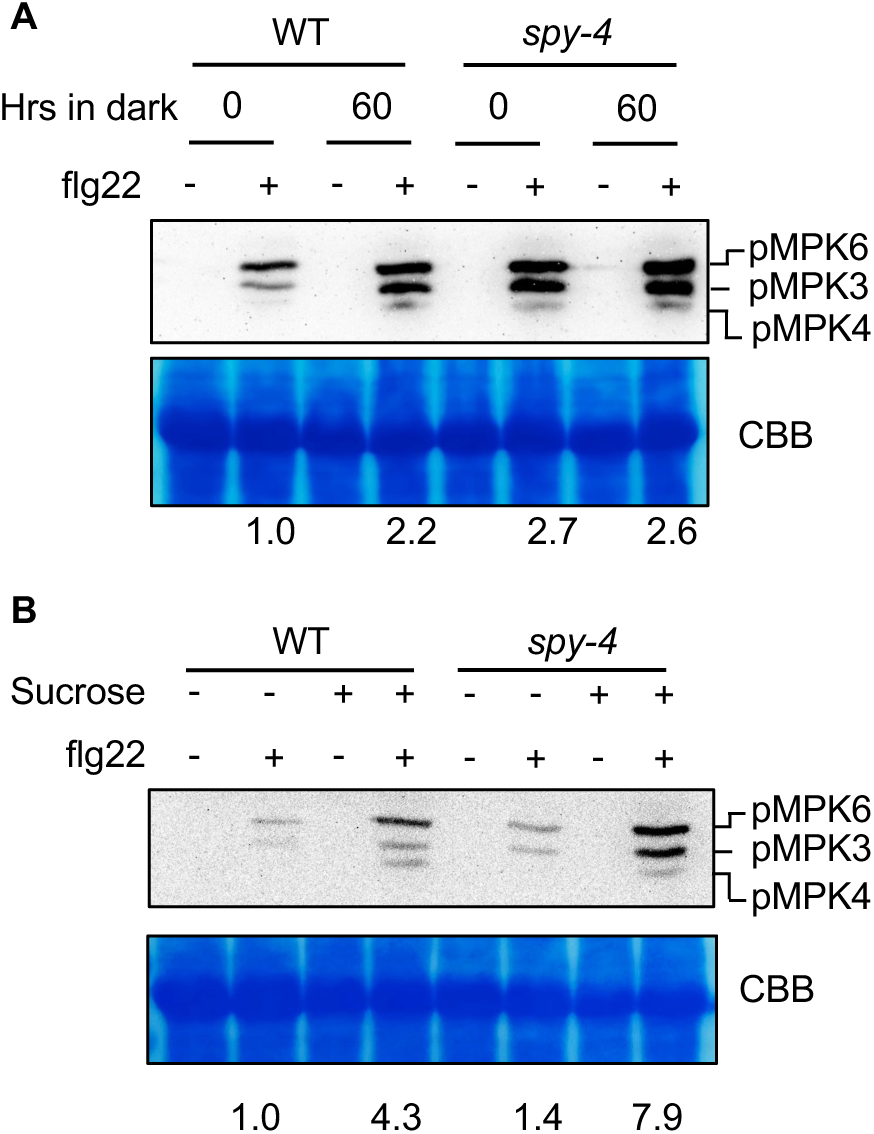
Sugar deprivation reduces O-fucosylation and enhances PTI response. Immunoblots showing MAPK phosphorylation level in WT and *spy-4* seedlings upon induction by 1 μM flg22 for 15 minutes. Seedlings were grown under light for 10 days and transferred into liquid medium with (B) or without (A) 1% sucrose for 60 hours. flg22 was added at the end of starvation treatment to induce MAPK phosphorylation. CBB: Coomassie Brilliant Blue staining of the gel blot.

**Figure S9.**
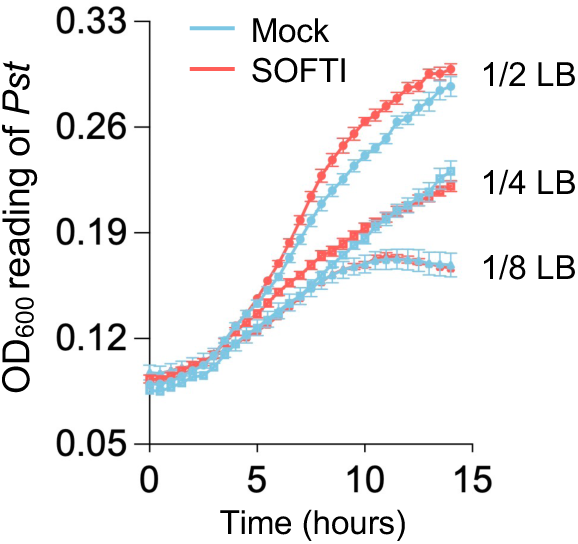
SOFTI does not inhibit *Pst* growth in liquid culture. Real-time recording of *Pst* DC3000 growth (Optical Density at 600 nm) in different dilutions of Lysogeny Broth (LB) medium containing 0 or 30 µM SOFTI over time (0-15 hours).

**Figure S10.**
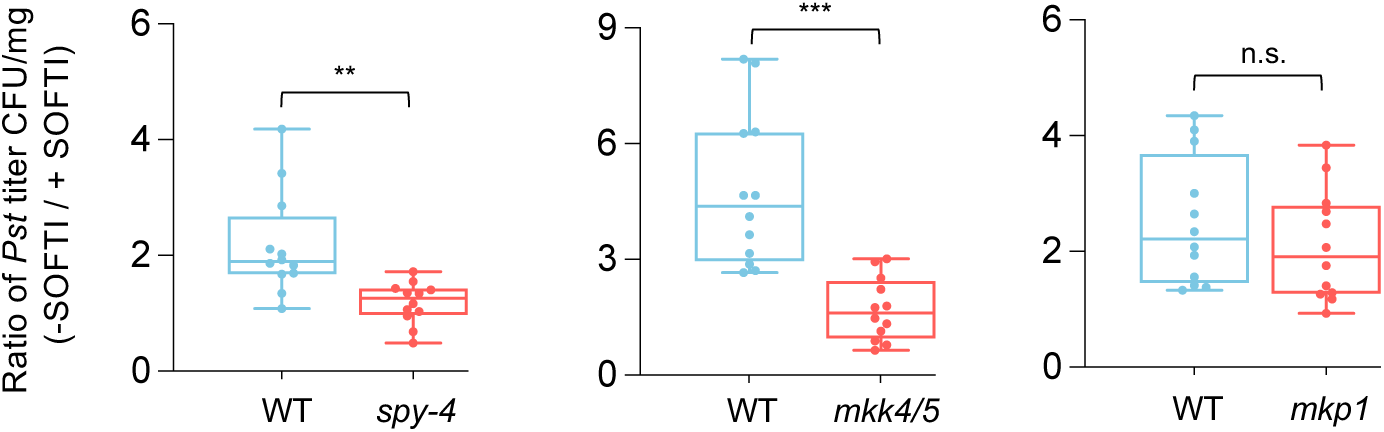
SOFTI enhancement of immunity depends on SPY and MKK4/5 but not MKP1. Ratio between *Pst* DC3000 titer (CFU/mg) of mock (-SOFTI) and SOFTI-treated (+SOFTI) seedlings described in Figure 6A-C. *** indicates *p* < 0.001, ** indicates p < 0.01, n.s. indicates *p* > 0.05 calculated with an unpaired Welch’s t test.

**Table S1. Oligonucleotides used in this study.**

**Table S2. Table S2 normalized reads for flg22-core responsive genes.**

## STAR methods

## KEY RESOURCES TABLE

| REAGENT or RESOURCE | SOURCE | IDENTIFIER |
| --- | --- | --- |
| **Antibodies** |  |  |
| anti-GFP, mouse | Boster | HT801-02 |
| anti-GST, mouse | Santa Cruz<br>Biotechnology | K1020 |
| anti-His, rabbit | Cell Signaling Technology | 2365 |
| anti-phospho serine/threonine, mouse | Phosphosolution | PP2551 |
| anti-p44/42 MAPK, rabbit | Cell Signaling Technology | 9102 |
| anti-SPY, rabbit | PhytoAB | PHY1737S |
| anti-HA, mouse | Cell Signaling Technology | 2367 |
| anti-myc, mouse | Thermo Scientific | 88843 |
| anti-O-fucose-Serine | GlycoScientific | FOS |
| anti-O-fucose-Threonine | GlycoScientific | FOT |
| biotinylated Aleuria aurantia lectin | Vector Laboratories | ZF0305 |
| HRP-conjugated Streptavidin | Thermo Scientific | 21134 |
| Goat Anti-Mouse IgG-HRP Conjugate | Bio-rad | 1706516 |
| Goat Anti-Rabbit IgG-HRP Conjugate | Bio-rad | 1706515 |
| **Bacterial and virus strains** |  |  |
| *E. coli* DH5 | Lab stock | N/A |
| *E. coli* BL21 (DE3) | Lab stock | N/A |
| *E. coli* Rosetta (DE3) | Lab stock | N/A |
| *Agrobacterium tumefaciens* GV3101 | Lab stock | N/A |
| **Chemicals, peptides, and recombinant proteins** |  |  |
| DMSO | Sigma-Aldrich | 472301 |
| SOFTI | MedChemExpress | HY-13465 |
| Hydrogen peroxide | Sigma-Aldrich | 216763 |
| ATP disodium salt trihydrate | USB Corporation | 117714 |
| GDP-fucose | Sigma-Aldrich | G4401 |
| UDP-GlcNAc | Sigma-Aldrich | U4375 |
| flg22 | MedChemExpress | HY-P1568 |
| pep1 | Protein and Nucleic Acid Facility at Stanford University | pan.stanford.edu |
| elf18 | Protein and Nucleic Acid Facility at Stanford University | pan.stanford.edu |
| RALF1 | Protein and Nucleic Acid Facility at Stanford University | pan.stanford.edu |
| chitin | Jeon et al.<sup>51</sup> | N/A |
| Triton X-100 | VWR life science | 0694-1L |
| Tween-20 | Promega | 0000364129 |
| NP-40 | Thermo Scientific | 85124 |
| Silwet L-77 | Phyto Tech Labs | S7777 |
| Pierce phosphatase inhibitor | Thermo Scientific | A32957 |
| Protease Inhibitor Cocktail | Thermo Scientific | 3214605 |
| PMSF | Sigma-Aldrich | P7626 |
| IPTG | MedChemExpress | HY-15921 |
| DTT | Goldbio | DTT10 |
| Imidazole | Sigma-Aldrich | I-2399 |
| L-Glutathione reduced | Sigma-Aldrich | G6629 |
| Luminol | Sigma-Aldrich | A8511 |
| Peroxidase from horseradish | Sigma-Aldrich | P8375 |
| Phos-tag Acrylamide AAL-107 | Fujifilm | 304-93521 |
| Zinc Chloride | Sigma-Aldrich | 211273 |
| Sodium metabisulfite | Thermo Scientific | 10231390 |
| Na<sub>2</sub>HPO<sub>4</sub> • 7H<sub>2</sub>O | Sigma-Aldrich | S9390 |
| KH<sub>2</sub>PO<sub>4</sub> • H<sub>2</sub>O | Thermo Scientific | 447670010 |
| Sodium Chloride | Macron | 7851-06 |
| Potassium Chloride | Sigma-Aldrich | P3911 |
| Magnesium Chloride | Sigma-Aldrich |  |
| SDS | Sigma-Aldrich | L3771 |
| Sucrose | VWR life science | M117 |
| Phytoblend | Caisson | PTP01 |
| Tris | VWR life science | M151 |
| Glycine | Sigma-Aldrich | G7126 |
| Protogel | National diagnostics | EC-890 |
| Ammonium persulfate | Research products international | 227004-246150 |
| PVDF membrane | Thermo Scientific | 88518 |
| TEMED | Sigma-Aldrich | 110732 |
| WesternBright Quantum kit | Advansta | 241118-34 |
| PureLink RNA Mini Kit | Invitrogen | 12183025 |
| All-in-One Ultra RT SuperMix | Vazyme | R433 |
| SupRealQ SYBR green mix | Vazyme | Q713 |
| GFP-Trap® Magnetic Agarose | ChromoTek | gtma |
| Pierce glutathione agarose resins | Thermo Scientific | 16101 |
| Superdex 200 column | Cytiva | 28990944 |
| anti-His biosensor | Gatorbio | 160009 |
| anti-GST biosensor | Gatorbio | 160042 |
| Zeba Spin Desalting Columns | Thermo Scientific | 89882 |
| Gibson assembly master mix | New England Biolabs | E2611L |
| PENTER/D-TOPO cloning kit | Thermo Scientific | 450218 |
| Q5 Site-Directed Mutagenesis Kit | New England Biolabs | E0552S |
| Phusion DNA polymerase | Thermo Scientific | F530S |
| Gateway LR Clonase II Enzyme mix | Thermo Scientific | 11791020 |
| **Experimental models:**<br>**Organisms/strains** |  |  |
| Col-0 | Widely distributed | N/A |
| *spy-4* | Aizezi et al.<sup>52</sup> | N/A |
| *spy-23* | Aizezi et al.<sup>52</sup> | N/A |
| *pSPY::SPY-GFP/spy-23* | Aizezi et al.<sup>52</sup> | N/A |
| *sec-2* | Bi et al.<sup>53</sup> | N/A |
| *sec-5* | Bi et al.<sup>53</sup> | N/A |
| *mkk4-18 mkk5-18* | Zhao et al.<sup>54</sup> | N/A |
| *mkp1* | Jiang et al.<sup>55</sup> | N/A |
| *MKK5-GFP* | Zhao et al.<sup>54</sup> | N/A |
| *MKK5-GFP/spy-4* | This paper | N/A |
| *MKK5-GFP/sec-2* | This paper | N/A |
| *fls2* | Lu et al.<sup>56</sup> | Salk_141277 |
| *BIK1::BIK1-HA* | Park et al.<sup>7</sup> | N/A |
| *UBQ10::GCaMP6* | Ast et al.<sup>57</sup> | N/A |
| *Solanum lycopersicum* cultivar VF36 | Widely distributed | N/A |
| **Oligonucleotides** |  |  |
| Primers are listed in Table S1 |  |  |
| **Recombinant DNA** |  |  |
| pDEST15-MKK4 K108R | This paper | N/A |
| pDEST15-MKK5 K99R | This paper | N/A |
| pCAMBIA1390-MKK5 | This paper | N/A |
| pEarleyGate 101-SPY | This paper | N/A |
| pSUMO-3TPR-SPY | Aizezi et al.<sup>52</sup> | N/A |
| pTrc-5TPR-SEC | Zentella et al.<sup>27</sup> | N/A |
| pSUMO-MEKK1 CA | This paper | N/A |
| **Software and algorithms** |  |  |
| ImageJ | NIH | https://imagej.nih.gov/ij/ |
| R version 4.5.1 | R Core Team | R: The R Project for Statistical Computing |
| Prism 10.3.0 | Graphpad | www.graphpad.com/guides/prism |
| R studio | Posit | https://posit.co/download/rstudio-desktop/ |

## EXPERIMENTAL MODEL AND STUDY PARTICIPANT DETAILS

### Plant material, growth conditions, molecular cloning, and statistical analysis

All Arabidopsis (*Arabidopsis thaliana*) plants used in this study are in Col-0 background. All seedlings were grown in long-day conditions (16 hours of light and 8 hours of darkness) with a light intensity of 82 μmol m^−2^s^−1^ at 22°C, except the plants used for flg22-induced growth inhibition assay were grown in short-day conditions (8 hours of light and 16 hours of darkness). All seeds were surface sterilized by immersing in 70% ethanol for 12 minutes and washed with sterile water 3 times, then put onto ½ MS plates supplemented with 1% sucrose and 7 g/L Phytoblend (Caisson Labs PTP01). Adult plants for bacterial inoculation assay were grown in short-day conditions (10 hours of light and 14 hours of darkness) at 22°C in soil pots. The T-DNA mutant lines *spy-4*^52^, *spy-23*^52^*, sec-2*^53^*, sec-5*^53^, and transgenic lines *pSPY::SPY-GFP/spy-23*^52^, *MKK5-GFP* ^54^, *BIK1::BIK1-HA*^7^ have been previously described. *MKK5-GFP/spy-4* and *MKK5-GFP/sec-2* were generated by crossing *MKK5-GFP* with *spy-4* and *sec-2* mutants.

Tomato (*Solanum lycopersicum*) plants used in this study are cultivar VF36, grown in a greenhouse with a 16-hour light and 8-hour dark cycle at a temperature range of 25°C to 28°C.

For *in vitro* expression and purification of GST-MKK4/5 proteins, the sequences of MKK4 and MKK5 were amplified using Phusion DNA polymerase (Thermo Scientific) with Arabidopsis cDNA as template. The amplified fragments were cloned onto pENTER/D-TOPO (Thermo Scientific) with attB × attP site-specific recombination (BP) reaction and subsequently cloned into the pDEST15 vector through the attL × attR site-specific recombination (LR) reaction (Thermo Scientific). The pCAMBIA1390-MKK5 for tobacco infiltration was also generated by a similar approach. pDEST15-MKK4 K108R and pDEST15-MKK5 K99R mutants were generated with Q5 Site-Directed Mutagenesis Kit (New England Biolabs) with primers listed in Table S1.

For *in vitro* expression and purification of MEKK1-CA protein, the truncated sequence of MEKK1 (326-592) was amplified using Phusion DNA polymerase (Thermo Scientific) with Arabidopsis cDNA as template. The amplified sequence was cloned onto the pRSFDuet-SUMO vector using Gibson assembly master mix (New England Biolabs).

All statistical analyses were performed by one-way ANOVA with Tukey’s multiple comparisons test to compare the mean of each group with every other group. **** indicates p < 0.0001, *** indicates p < 0.001, ** indicates p < 0.01, * indicates p < 0.05, and n.s. indicates p > 0.05.

## METHOD DETAILS

### Protein extraction and immunoblot analysis

Plant samples were ground into fine powder with liquid nitrogen. Proteins were extracted using 2×SDS loading buffer (60 mM Tris–HCl, pH 6.8, 25% v/v glycerol, 2% SDS, 100 µM Dithiothreitol, and 0.1% bromophenol blue) and boiled at 65 °C for 10 mins. The samples were then loaded onto 10% SDS-polyacrylamide gels and transferred onto polyvinylidene difluoride (PVDF) membrane. The membrane is blocked with 3% BSA dissolved in PBST solution (2 mM KH_2_PO_4_, 10 mM Na_2_HPO_4_, 137 mM NaCl, 2.7 mM KCl, Tween-20 added at 0.1% v/v) overnight on a 4 °C shaker and further probed with anti-GFP (Transgene, Q20329, 1:2000 dilution), anti-GST (Santa Cruz Biotechnology #K1020, 1:2000 dilution), anti-His (Cell Signaling Technology #2365, 1:2000 dilution), anti-phosphoserine/threonine (Phosphosolution #PP2551, 1:2000 dilution), anti-p44/42 MAPK (Cell Signaling Technology #9102, 1:2000 dilution), anti-SPY (PhytoAB PHY1737S, 1:1000 dilution), or biotinylated Aleuria aurantia lectin (AAL, Vector Laboratories ZF0305, 1:5000 dilution) dissolved in 3% BSA for 1 hour. After washing with PBST for 3 times at 6-minute intervals, the membrane is probed with anti-mouse or anti-rabbit secondary antibodies (BIO-RAD STAR132/STAR121, 1:5000 dilution) or Pierce HRP-conjugated Streptavidin (Thermo Scientific, 1:10,000 dilution) for 45 minutes and then washed with PBST for 3 times at 6-minute intervals. A chemiluminescence assay was used for band visualization using the WesternBright Quantum kit (Advansta).

### Phos-tag SDS PAGE analysis of phosphorylated proteins

The 10% separating gel for Phos-tag experiment was made as follows: mix 2085 µL of 30% acrylamide-bis-acrylamide (29:1 ratio, National diagnostics), 2185 µL of 1 M Tris (pH = 6.8), 62.5 µL of 10% SDS, 93.8 µL of 5 mM Phos-tag (Fujifilm), 93.8 µL of zinc chloride, 25 µL of 10% APS, 5 µL of TEMED, and 1715 µL of double distilled water.

The 4% stacking gel was made as follows: mix 417.5 µL of 30% acrylamide-bis-acrylamide (29:1 ratio, National diagnostics), 1093 µL of 1 M Tris (pH = 6.8), 31.3 µL of 10% SDS, 25 µL of 10% APS, 5 µL of TEMED, and 1570 µL of double-distilled water.

Protein sample was prepared in the same way as a regular SDS-PAGE as described previously, and gel electrophoresis was done with 1x MOPS running buffer: mix 12.1 g of Tris, 20.9 g of MOPS, 1 g of SDS, and 0.95 g of sodium metabisulfite, pH = 7.8. The entire apparatus was dipped in ice water bath to keep the temperature cool during the run. The gel was then immersed in 1x Transfer buffer (mix 3 g Tris, 14.4 g Glycine, 200 mL methanol, and 800 mL of double-distilled water) containing 1 mM EDTA for 10 minutes with shaking, then immersed in 1x transfer buffer without EDTA for 20 minutes. Proteins were then transferred to a PVDF membrane for immunoblot analysis.

### Real-time recording of calcium reporter signal

Roots of 10-day-old seedlings of the *pUBQ10::GCaMP6* reporter line were excised, and 3 roots were transferred into one well from a 96-well black plate containing 100 µL of sterile water. The roots were then put under bench light for 6 hours. Then, 100 µL of 2 µM flg22 or sterile water was added into each well, and the plate was immediately put into a plate reader (Tecan infinite M1000 pro) and scanned at excitation: 488 nm and emission: 509 nm for 50 minutes.

### Reactive Oxygen Species (ROS) burst assay

Leaf discs (4 mm diameter) from 4.5-week-old Arabidopsis plants (n = 12 leaf discs from different leaves) were incubated in sterile water in a 96-well plate (one leaf disc per well) for 16 hours. Before measurement, the water was removed from each well by vacuum, and 100 µL reaction mix containing 1 µM flg22, 20 µg/mL horseradish peroxidase, and 200 µM luminol (Sigma #123072) was added into each well of the plate. The plate was immediately measured in a Synergy H1 Microplate Reader (Biotek). Relative luminescence units (RLUs) are plotted against time.

### Pathogen infection and growth assay

To analyze the effect of pathogen infection on protein O-fucosylation, leaves of 4.5-week-old Arabidopsis plants were infiltrated with *Pseudomonas syringae* pathovar *tomato* (*Pst*) strain DC3000 (1 x 10^5^ colony-forming units per milliliter in sterilized 10 mM MgCl_2_) or 10 mM MgCl_2_ as a mock control. Plants were kept under a dome after infiltration to maintain humidity. Three days after inoculation, the leaves were flash-frozen in liquid nitrogen and further analyzed by nuclear protein extraction and immunoblot.

For pathogen growth assays, Arabidopsis seedlings were infected with *Pst* DC3000 according to a previously described protocol ^58^, with slight modifications. Arabidopsis seedlings were grown on ½ MS plates supplemented with 1% sucrose with or without the addition of 20 µM SOFTI for 7 days. *Pst* DC3000 was diluted to 1 x 10^6^ colony-forming units per milliliter with sterilized water containing 0.025% Silwet L-77. The bacterial inoculum was added to Arabidopsis plates to cover the seedlings for 2 minutes and then discarded. Three days after inoculation, seedlings were surface-sterilized, and the bacterial titers in the tissues were determined. Each group consisted of 12 biological replicates, and each replicate contained 3 seedlings.

For infection of tomato leaves, *Pst* DC3000 was diluted to 1 x 10^5^ colony-forming units per milliliter for pathogen growth assay and 2 x 10^4^ colony-forming units per milliliter for disease symptoms assay. Two leaflets of a pair on a leaf were pre-treated by infiltration of mock and 20 µM SOFTI solution a day before bacterial infection. Bacterial solutions supplemented with mock solvent DMSO or SOFTI were infiltrated into the two leaflets. The *Pst* titer was measured four days after inoculation, and the disease symptoms were recorded five days after inoculation.

### RNA extraction, sequencing analysis, and qRT-PCR

10-d-old Arabidopsis seedlings (Col-0, *spy-4*, and *sec-2*) were immersed in water for 16 hours and then treated with 100 nM flg22 peptide for 1 hour. The samples were then flash frozen and ground to fine powder in liquid nitrogen. Plant total RNA was extracted using a PureLink RNA Mini Kit (Invitrogen).

RNA sequencing was performed by Novogene to generate 150 bp paired-end reads with three biological repeats for each treatment. Raw RNA-seq reads were quality-checked with FastQC and aligned to the *Arabidopsis thaliana* reference genome, TAIR10^59^ using HISAT2^51^. Gene-level read counts were quantified with featureCounts^60^ based on the TAIR10 gene annotation (release 62). Differential gene expression analysis was performed using the DESeq2 package in R^61^. Genes with fewer than 20 reads across all samples were excluded. Raw counts were normalized using the median ratio method implemented in DESeq2, which estimates size factors to adjust for differences in sequencing depth among samples.

For data visualization, K-means clustering was performed on the normalized counts of a subset of 205 flg22-induced core immune genes. This gene set was defined as those upregulated at 10, 30, 90, and 180 minutes post-flg22 treatment from a broader list of 970 core immune response genes, as described by Bjornson et al.^62^

For qRT-PCR, the extracted RNA was reverse-transcribed with an All-in-One Ultra RT SuperMix (Vazyme). Quantitative real-time PCR was performed with SupRealQ SYBR green mix (Vazyme) with the primers listed in table S1 in a LightCycler 480 instrument (Roche).

### AAL affinity purification assay

The AAL affinity purification assay was performed with a previously described protocol with slight modifications.^32^ 100 mg of tissue from MKK5-GFP in Col-0 or *spy-4* backgrounds was homogenized with metal beads in liquid nitrogen. Five hundred microliters of extraction buffer (50 mM Tris-HCl, pH = 7.5, 50 mM NaCl, 10% v/v glycerol, 0.25% v/v Triton-X 100, 0.25% v/v NP40, 1 mM PMSF, 1× Halt protease inhibitor cocktail [Thermo Scientific], 1× Pierce phosphatase inhibitor [Thermo Scientific]) was added to each sample. The nucleus was lysed by 1 minute of ultrasonication, and the samples were then centrifuged at 14,000 rpm for 10 minutes. The supernatant was incubated with 30 µL of AAL-conjugated agarose beads (Vector Labs, # AL-1393-2) for 1.5 hours at 4°C with end-to-end rotation and then washed with lysis buffer 5 times and boiled with 30 µL 2xSDS loading buffer for 10 minutes at 65 °C and further analyzed by immunoblot analysis.

### Protein Immunoprecipitation

One hundred milligrams of Arabidopsis tissue or 250 milligrams of *Nicotiana benthamiana* was homogenized with metal beads in liquid nitrogen and 200 microliter of ice-cold extraction buffer (50 mM Tris-HCl, pH = 7.5, 50 mM NaCl, 10% v/v glycerol, 0.25% v/v Triton-X 100, 0.25% v/v NP40, 1 mM PMSF, 1× Halt protease inhibitor cocktail [Thermo Scientific], 1× Pierce phosphatase inhibitor [Thermo Scientific]) was added to each sample. Each sample was ultrasonicated for 1 minute to break the nucleus and centrifuged at 14,000 rpm for 10 minutes. Twenty µL of supernatant was collected as input and boiled with 20 µL of 2xSDS loading buffer. The supernatant was then incubated with 20 µL of anti-GFP magnetic beads (ChromoTek GFP-Trap® Magnetic Agarose) for 1 hour in 4°C with end-to-end rotation. The magnetic beads were then collected with a magnetic stand and washed with extraction buffer 5 times and boiled with 30 µL of 2xSDS loading buffer for 10 minutes at 65 °C and further analyzed by immunoblot analysis.

### Recombinant protein purification

The cDNA sequences of SPY (residues 326-914), SEC (residues 356-977), MEKK1 (residues 326-592), and PCK1 were cloned into RSFDuet-SUMO vector with an N-terminal 6xHis-SUMO tag. Those of MKK4 (K99R) and MKK5 (K108R) were cloned into pDEST15 vector containing an N-terminal GST tag. The point mutations were generated via site-directed mutagenesis. All constructs were transformed into Rosetta DE3 *E. coli* strains for protein expression. The strains were grown until OD_600_ = 0.8, and 300 µM isopropyl-β-D-thiogalactoside (IPTG) was added to induce protein expression at 16°C for 6xHis-SUMO tagged proteins and 22°C for GST-tagged proteins overnight. Cell pellets were then collected, resuspended in 1×PBS (2.7 mM KCl, 2 mM KH_2_PO4, 137 mM NaCl, 10 mM Na_2_HPO4, 1mM PMSF, and 1× Halt protease inhibitor cocktail [Thermo Scientific], pH = 7.4), and lysed with ultrasonication. The proteins were then affinity-purified with Ni^2+^ columns (Invitrogen, R901-01) or Pierce™ glutathione agarose resins (Thermo Scientific). The eluted proteins were further purified with gel filtration chromatography using a Superdex 200 column (GE Healthcare) on an Akta purifier (Cytiva). The final protein product was concentrated to 1mg/mL and stored in 1×PBS at −70 °C.

### In vitro GST pull-down assay

Five µg of GST, GST-MKK4 (K108R), or GST-MKK5 (K99R) recombinant proteins were immobilized onto 30 µL Pierce™ glutathione agarose resins (Thermo Scientific) by incubating at 4°C for 4 hours. After 5 washes with 10 mL 1xPBST (0.1% Tween-20, v/v), the beads were incubated with 5 µg of 6xHis-SUMO tagged SPY, SEC, and PCK1 proteins for 1 hour at room temperature. The beads were then washed again with 1xPBST for 5 times, each time with end-to-end rotation for 3 minutes, and eventually boiled in 50 µL 2xSDS loading buffer at 95 °C for 10 minutes and further analyzed by immunoblot.

### Bio-layer interferometry (BLI) assay

BLI assay was performed with a previously described protocol with slight modifications ^52^. In brief, bait proteins (final concentration 20 µg/mL) were immobilized onto anti-His (Gator bio #160009) or anti-GST biosensors (Gator bio, #160042) using a Gator-plus instrument (Gator Bio). The sensors were then dipped into analyte proteins at indicated concentrations; the AAL concentration was fixed at 20 µg/mL. The dissociation constant (K_d_) was calculated by steady-state fitting of the response unit to serial dilutions of the analyte protein.

### *In vitro O*-glycosylation and phosphorylation assays

Five µg of recombinant GST-MKK5 (K99R) protein was incubated with five µg of recombinant 3TPR-SPY and/or 5TPR-SEC in the presence of 1 mM GDP-fucose and/or UDP-GlcNAc at room temperature overnight or as indicated in the reaction buffer (1xPBS, pH = 7.3, 5 mM MgCl_2_). The reaction product was desalted with Zeba Spin Desalting Columns (Thermo Scientific) to remove the GDP and UDP. Five µg of recombinant MEKK1-CA protein with 1 mM ATP was added to the reaction product and incubated at room temperature for 1 hour. GST-MKK5 proteins were then immobilized onto 30 µL Pierce™ glutathione agarose resins (Thermo Scientific) by incubating at room temperature for 1 hour. The resin was then washed with 1xPBST containing 300 mM NaCl for 5 times with end-to-end rotation, 3 minutes each. The GST-MKK5 protein was eluted by adding 2xSDS loading buffer and boiling at 65°C for 10 minutes. The samples were further analyzed by immunoblot analysis.

### Nuclear protein fractionation

At least 100 mg of Arabidopsis seedlings were flash frozen and ground in liquid nitrogen to fine powder. Then, 500 µL ice-cold extraction buffer 1 (0.4 M sucrose, 10 mM Tris-Cl pH = 8.0, 5 mM dithiothreitol, 0.1 mM PMSF, and cocktail protease inhibitors) was added. The extract was filtered with a 70 µM cell strainer mounted on a 50 mL conical tube and centrifuged at 800 x *g* for 3 minutes at 4°C. The pellet was resuspended with supernatant and transferred into a 1.5 mL centrifuge tube, then centrifuged at 700 x *g* for 5 minutes at 4°C. The pellet was resuspended in 1 mL of buffer 2 (0.25 M sucrose, 10 mM Tris-Cl, pH = 8.0, 5 mM dithiothreitol, 10 mM MgCl_2_, 1% Triton X-100 v/v, 0.1 mM PMSF, and cocktail protease inhibitors). The sample was centrifuged at 700 x *g* for 5 minutes at 4°C. The supernatant was removed, and pellet was resuspended in 500 µL of buffer 3 (1.7 M sucrose, 10 mM Tris-Cl pH = 8.0, 5 mM dithiothreitol, 2 mM MgCl_2_, 0.15% Triton X-100 v/v, 0.1 mM PMSF, and cocktail protease inhibitors) and layered on top of another 500 µL of buffer 3, then centrifuged at 15, 000 *g* for 10 minutes at 4°C. The nuclear pellet was then resuspended in 50 µL 2xSDS loading buffer and boiled in 95°C for 5 minutes. The samples were then analyzed by immunoblot analysis.

### Mass spectrometry analysis

After Colloidal Blue staining, the MKK4 and MKK5 protein bands were excised and subjected to in-gel digestion with trypsin. The resulting peptide samples were desalted using C18 ZipTips (Millipore) and analyzed on a Q-Exactive HF hybrid quadrupole-Orbitrap mass spectrometer (Thermo Fisher) equipped with an Easy-nLC 1200 UHPLC liquid chromatography system (Thermo Fisher). The MS data were searched using Protein Prospector against the Arabidopsis Information Resource (TAIR10) database. Reverse protein sequences (35,386 entries) were concatenated to the database to allow the estimation of false discovery rate (FDR). Carbamidomethylation of cysteine was set as a fixed modification. Oxidation of methionine, N-terminal acetylation, *O*-GlcNAcylation of serine/threonine, neutral loss of *O*-GlcNAcylation, *O*-fucosylation of serine/threonine, and neutral loss of *O*-fucosylation were set as variable modifications. The precursor ion mass tolerance was set to 10 ppm. and the fragment ion mass tolerance to 20 ppm Peptide and protein FDRs were set as 0.01 and 0.05, respectively.

